# The complexity of multiple CRISPR arrays in strains with (co-occurring) CRISPR systems

**DOI:** 10.1101/2025.05.05.651427

**Authors:** Axel Fehrenbach, Alexander Mitrofanov, Rolf Backofen, Franz Baumdicker

## Abstract

CRISPR and their associated Cas proteins provide adaptive immunity in prokaryotes, protecting against invading genetic elements. These systems are categorized into types and are highly diverse. Genomes often harbor multiple CRISPR arrays varying in length and distance from Cas loci. However, the ecological roles of multiple CRISPR arrays and their interactions with multiple Cas loci remain poorly understood.

We present a comprehensive analysis of CRISPR systems that uncovers variation between diverse Cas types regarding the occurrence of multiple arrays, the distribution of their lengths and positions relative to Cas loci, and the diversity of their repeat sequences. Some types tend to occur as the sole Cas locus present in the genome, but typically have two or more associated arrays, especially for types I-E and I-F. Multiple Cas types are also common, with some systems showing a preference for specific co-occurrence. Distinct array distributions and orientations around Cas loci indicate substantial differences in functionality and transcriptional behavior among Cas types.

Our analysis suggests that arrays with identical repeats in the same genome acquire new spacers at comparable rates, irrespective of their proximity to the Cas locus. Furthermore, repeat similarities indicate that arrays of systems that often co-occur with other systems tend to have more diverse repeats than those mostly appearing alongside solitary systems. Our results indicate that co-occurring Cas type pairs might not only collaborate in spacer acquisition but also maintain independent and complementary functions and that CRISPR systems distribute their defensive spacer repertoire equally across multiple CRISPR arrays.

## Introduction

CRISPR systems provide adaptive immunity in bacteria and archaea by incorporating short DNA fragments from invading genetic elements, such as phages and plasmids, into CRISPR arrays. These fragments, so called spacers, enable sequence-specific recognition and degradation of future invaders. CRISPR-Cas plays an important role in shaping microbial evolution and community dynamics [1]. The distribution of CRISPR systems and, in particular, the variability of their CRISPR arrays offer insights into the defense strategies and evolutionary pressures exerted by phages and mobile genetic elements [2, 3].

Prokaryotic genomes encode a remarkable variety of CRISPR systems that differ in their molecular components and organization and provide distinct functions [4, 5]. CRISPR systems are widespread, can be encoded on the chromosome as well as plasmids [6], but exhibit a patchy distribution across bacterial genomes, which indicates that CRISPR systems also carry a cost [7]. Multiple types of CRISPR systems can coexist within a single strain [8, 9] which can lead to interactions between the systems and the associated arrays, such as spacer acquisition in trans [10–15].

In addition to the diversity of types and variety of Cas proteins, the variability of the associated CRISPR arrays are of interest. The spacer arrays provide a partial track record of spacer acquisitions that occurred, e.g. due to phage infection attempts and reflect the defensive capabilities of a cell. Since the frequency of transcription and the degree of immunity conferred by the spacers are influenced by the length of the array and the positions of the spacers [14, 16, 17], the composition of CRISPR arrays can provide insights into the arms race between bacteria and phages [3]. Arrays vary in their configuration; they may be located near the Cas genes or dispersed throughout the genome, and can differ in their length (i.e. the number of spacers), leader and repeat uniformity. The repeat sequences, which intersperse with spacer sequences within CRISPR arrays, are known to be type-specific to CRISPRs [9, 18]. The leader sequence plays an important role in regulating spacer integration and expression. In general, the leader sequence is located upstream of the CRISPR array and has been found to be very diverse, containing motifs that are type-but also taxonomy-specific [18, 19]. Together this variability suggests that besides the repeats and leaders, the structure and positioning of CRISPR arrays have important implications for the functional efficiency and adaptability of CRISPR systems.

When multiple CRISPR arrays exist within a bacterial genome, it remains unclear if and if so which arrays are preferentially used for new spacer acquisitions. One key factor may be their genomic proximity to Cas genes, as closer arrays might be more actively transcribed, while distant arrays could serve as archival records. Additionally, the presence of multiple arrays raises the question of whether they serve distinct functional roles or provide redundancy. While previous studies and simulations have explored spacer acquisition dynamics and array length distributions [8, 9, 20], a large-scale comparative analysis is still missing. Similarly, although the interaction between multiple CRISPR-Cas systems has been studied [21], it is not yet fully understood. If different systems coexist, do their arrays exhibit more similar repeats, potentially allowing for the cross-utilization of Cas proteins, are certain systems incompatible with each other, or do they function independently? Understanding these relationships can shed light on the evolutionary pressures shaping the diversity of CRISPR arrays and systems.

Here, we systematically explore a database with thousands of prokaryotic genomes, their CRISPR systems and CRISPR arrays. We utilize the relationship between mean spacer acquisition rates and array lengths to investigate the relative activity of multiple CRISPR arrays and how the array position relative to Cas genes and the repeat sequence influences spacer acquisition. We also consider how multiple CRISPR systems within a genome interact, and find that all these patterns differ substantially between Cas types. We hope that the comprehensive analysis here provides the basis for a better understanding of these differences and can shed light on why certain array configurations are favored and how they impact bacterial immunity.

## Methods

### Dataset

We used a comprehensive dataset of CRISPR arrays obtained by extracting available whole genomes from GenBank (in November 2024) and running CRISPRidentify [22] to detect all high confidence (labeled by CRISPRidentify as “bona-fide” and “possible”) CRISPR arrays within each genome. Cas genes were identified and labeled by CRISPRCasidentifier [23]. Cas types are assigned according to the closest Cas locus, if no Cas was found within the genome they are assigned “no cas type” and called “orphans”. These orphan arrays have been excluded for all major analyzes and figures but appear in some supplemental figures and tables, as indicated.

Arrays are labeled as “close” to their Cas, if they are less than 10000 bp from their respective Cas locus. Array orientations were determined using CRISPRstrand [24]. We obtained a total of 47789 genomes with 82981 CRISPR arrays, 44760 distinct Cas loci and 19946 unique consensus repeats; see Supplementary Tables S1–S4 and Supplementary Fig. S1–S5 for additional statistics.

### Cas type co-occurrence analysis

We also considered the co-occurrence of Cas types relative to their total abundance. For each possible combination of Cas types, we compared the observed and expected numbers, where the expected number is given by the mean number of such genomes when the presence and absence of Cas types would be randomly and independently distributed across the dataset according to the observed frequency. We did this (i) for the observed and expected number of times a system occurs as the only system within a genome, and (ii) the observed and expected number of times a pair of CRISPR systems co-occurs. For example, if for a total of *n* genomes, one Cas type is present in *j* genomes and another type is present in *k*, not necessarily distinct, genomes, the expected number of genomes where both types co-occur would be *jkn*^−2^. The resulting ratios of observed and expected numbers are transformed into a fold-change and are depicted in Figure 1. The corresponding data and p-values are given in Supplementary Tables S5 and S6. A positive foldchange of −*x* indicates that the observed number is *x*-times more frequent than expected, whereas a negative fold-change of − *x* indicates that the observed number is *x*-fold smaller than expected.

**Fig. 1.**
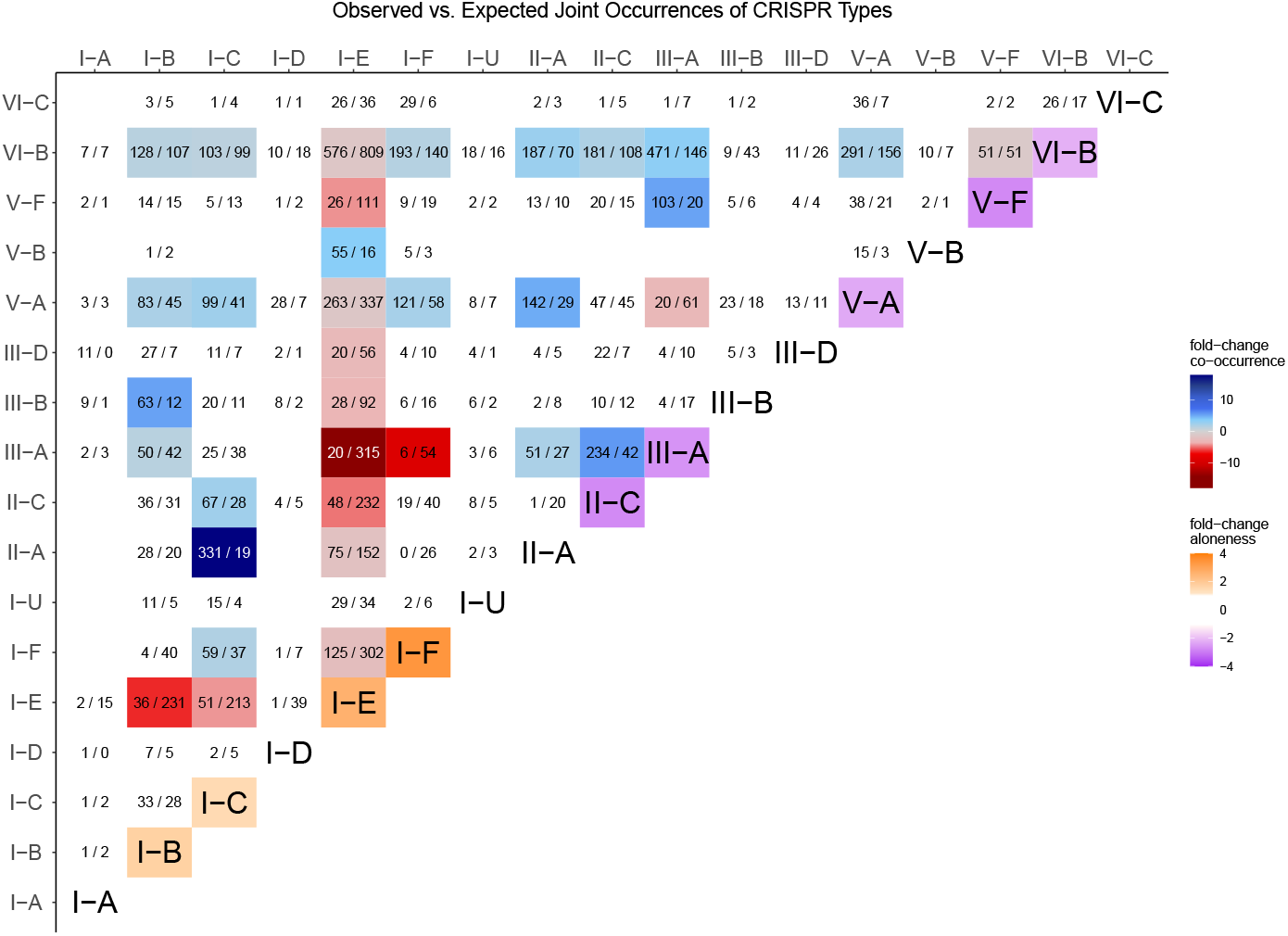
Single and Co-occurring CRISPR Types. For each pair of different Cas types we show the actually observed and the expected (if their occurrence would be reshuffled) number of joint occurrences (in the format observed/expected). For system-pairs where at least 50 genomes are observed or expected the color indicates the fold-change between the observed and the expected number. I.e. a -5 fold-change is shown in red and indicates that the number of observed co-occurrences is 5-fold smaller than expected if CRISPR systems would have been reshuffled, while blue colors indicate a more frequent co-occurrence. On the diagonal the color indicates a different fold-change. Namely, the tendency for each type to occur as the only CRISPR system (positive fold-change, orange) or to occur in combination with at least one other system (negative fold-change, purple). For example types I-E and I-F are alone 2.6-fold and 3.4-fold more often than expected and II-C occurs alone 1.7-fold less often than expected. On the diagonal only systems with at least 50 expected or observed solo ocurrences and with a significant fold-change (bionomial test p-value < 0.0001) got colored.

### Pairwise distances between consensus repeats

We compute any distances between repeats solely for the respective consensus repeats of the arrays. For any two consensus repeats we compute the Levenshtein distance between the two. These distances are normalized by the maximum length of consensus repeats within the genome for Supplementary Fig. S13 or globally for Figure 4, respectively.

### Visualization of repeat distances with multidimensional scaling (MDS)

We computed the pairwise Levenshtein distances for all consensus repeats for all arrays in the dataset as described above. Then, we performed metric MDS, using the obtained distance matrix as dissimilarity measure, to obtain a projection into two-dimensional space using the MDS function of the python package scikit-learn (v. 1.3.2). This MDS function uses the metric SMACOF algorithm and minimizes the stress (residual sum of squares of the original distances and the distances in the 2-d MDS space). The projection of each array’s consensus repeat into the MDS space is shown in Figures 4 and 5. All subfigures show subsets of arrays projected using the *same* MDS projection for greater visual clarity.

In addition to the scatterplots of the repeats projected into the two-dimensional MDS space, we provide in Figure 5 kernel density estimates with Gaussian kernels of the marginal distributions for both MDS coordinates and the indicated (colored) subsets, respectively. To compute the kernel density estimates we use the kdeplot function of the python package seaborn (v. 0.12.2) with default parameters, which relies on SciPy (v. 1.9.1) for the Gaussian kernel density estimates.

### Estimation of comparative insertion rate weights

To compute the insertion rate weights (Figure 3) we assume an independent deletion model (IDM) as described in [2] which evolves by independent acquisitions and deletions of spacers with insertion rate *θ* and deletion rate *ρ*.

In the IDM the array length is Poisson distributed with mean *θ/ρ* and thus the average length *n* is given by *θ/ρ*. As found in [2], deletion rates were found to be independent of Cas type and likely caused by processes, e.g. replication misalignment, which should be consistent for different arrays within a single genome. Thus, we assume the deletion rate to be constant and the insertion rate is likely the main driver of differing array lengths of arrays within the same genome.

Therefore, for a genome with *k* arrays we can roughly estimate the insertion rate of array *i* ∈ {1, …, *k*} as *θ*_*i*_ = *n*_*i*_*ρ*, where *n*_*i*_ is the length of the array. Furthermore, we define the total insertion rate for all arrays in the genome as *θ* = *θ*_1_ + … + *θ*_*k*_ for arrays *i* ∈ {1, …, *k*} within one genome with *k* spacer arrays. To compare the acquisition behavior of CRISPR arrays within the same genome we compute what we call (relative) *insertion rate weights* given by *w*_*i*_ = *θ*_*i*_*/θ*. Clearly, by definition of *θ* and *θ*_*i*_, and due to the constant deletion rate, *w*_*i*_ is solely determined by the proportions of array lengths, i.e. we find *w*_*i*_ = *n*_*i*_*/n*, where *n* = *n*_1_ + … + *n*_*k*_ is the combined total length of all arrays.

To obtain the expected neutral distributions of insertion rate weights in Figure 3 we proceeded as follows.

In a neutral setting, where all arrays are equally likely to acquire new spacers, for a genome with *k* arrays, any new spacer is inserted into any of the arrays with equal probability (1*/k*). Thus, given a total of *n* spacers, the array lengths of the *k* arrays would be distributed according to a multinomial distribution with *n* number of trials, *k* distinct events and equal probabilities *p*_*i*_ = 1*/k* for *i* ∈ {1, … *k*} . We therefore sampled using the following approach: for each datapoint, i.e. genome, with *k* arrays and the total number of spacers *n* we collected a single sample from the corresponding multinomial distribution and divided by *n* to obtain the insertion rate weights. The resulting distribution thus has the same total number of samples as the real data analyzed.

## Results & Discussion

### Diversity and co-occurrence of Cas types

We found that although the distribution of Cas types is dominated by a few abundant types such as I-E, I-F, III-A, V-A, VI-B and a substantial amount of orphan arrays (where no Cas locus was found in the genome), there is substantial diversity to explore (see Supplementary Tables S1–S3 and Supplementary Fig. S1, S2).

Moreover, although most genomes carry less than 7 distinct Cas loci with a respective CRISPR array (Supplementary Fig. S3) of less than 4 distinct types (Supplementary Fig. S4), a substantial number of genomes carry at least two different Cas types. Notably, certain combinations of Cas types are much more common than others, suggesting a potential interaction or synergy between some types [25]. For example, based on the considered dataset this might be the case for II-A and I-C, as well as II-C and III-A, and also III-A and V-F (see Figure 1) as those systems often occur together.

On the other hand, some systems, including I-E and I-F, tend to occur as the only CRISPR system more often than expected, which could indicate that these systems either provide already sufficient defense or that the presence of other systems interferes with their function.

In this context, it is also important to consider horizontal gene transfer (HGT) as one major vector in the propagation of CRISPR systems. Often, CRISPR systems are transferred via plasmids. There is clear evidence of CRISPR systems on (larger) plasmids, most of which are of type I and IV [6, 26]. Type IV systems, represented by only a few instances in our dataset, are predominantly plasmid-encoded [6]. Thus, HGT, indirectly, plays a role in shaping the distribution of CRISPR systems and their co-occurrence. However, since we are considering a comprehensive, cross-sectional dataset, we expect our findings to be robust against noise introduced by HGT.

Moreover, it is believed that HGT is (often) inhibited by already existing CRISPR systems in the receiving genome. Bernheim et al. [21] propose that systems that are incompatible with a functioning system in the cell would lead to the rapid deletion of one of the systems. For example, in *Escherichia coli* most strains contain a functional I-E system and at least one type I-F array without Cas genes where the majority of spacers within the I-F array target absent type I-F Cas genes [27, 28]. Bernheim et al. [21] argue that this leads to immediate degration of incoming I-F Cas genes and thus to the inability of horizontal transfer of functional type I-F systems into *E. coli*. Plasmid-born arrays have been found to frequently carry spacers that target (other) plasmids [6]. Furthermore, a recent study [29] showed that indeed, the exchange of mobile genetic elements, and thus HGT, is significantly inhibited by CRISPR systems. However, they found this effect to be highly variable between species.

While other studies considered the co-occurrence of Cas types [8, 9, 21], here we also consider the number and length of the associated spacer arrays and the similarity of repeat sequences to obtain a more comprehensive perspective.

### Cas loci often support multiple arrays

Clearly, the number of arrays tends to exceed the number of Cas loci, indicating that many Cas loci can and do serve multiple arrays (Supplementary Fig. S5). Although many Cas loci have only one array, we found that many Cas loci have multiple close CRISPR arrays, as there is an overall average of 1.48 arrays per Cas locus within the dataset (Supplementary Table S1 and Supplementary Fig. S6). Larger amounts of around 5 arrays do occur quite often and larger numbers (up to around 10) are rare, but do appear (with a maximum of 23 for one V-A locus, see Supplementary Table S1).

Particular outliers are types I-E, I-F and III-A since they have a median of two arrays which we will discuss further below. On the other hand, Type II-A strongly favors only a single array (average 1.07). Similar observations have been made before based on the signature cas genes for type I and II [8]. Our analysis shows that all Cas types have at least some Cas loci with multiple arrays. These arrays originate from duplication (of fragments of an array), multiple acquisitions of arrays (via HGT) and splitting of arrays into multiples due to mutations or insertions, and subsequently change in spacer content.

Duplications of arrays could be a major contributor for the frequency of same consensus repeat arrays. Recently duplicated arrays would have the same consensus repeat and likely share large subsets, or possibly all, of the spacers. Although CRISPR arrays diverge in their spacer composition relatively quickly, the last spacer at the leader distal end is the least likely to be deleted and would therefore be more likely shared by arrays with somewhat distant ancestry [2].

To get a sense of the frequency of recent duplications, we compared the repeats and spacer content of all arrays and Cas loci and report the statistics in Supplementary Tables S1 and S2. We found that similar consensus repeats are frequent. Of all arrays, 25.0% co-occur in a genome with at least one additional array with identical consensus repeat and 19.3% of the Cas loci have multiple associated arrays with the same consensus repeats. A substantial part of those arrays (11.9%) have at least some spacer overlap. However, only relatively few arrays (1.2%) have identical spacers. Thus, recent complete duplication or complete absence of spacer evolution is rare in our data set. A higher percentage (4.1%) have the same last spacer and thus more clearly indicate common ancestry. These values vary significantly between Cas types (Supplementary Table S2).

In general, the relatively high frequency of multiple CRISPR arrays suggests that, at least for some types, there is a beneficial fitness effect [21], such as robustness due to redundancy or specialization of arrays on specific targets [8, 21]. However, definitive empirical evidence for these theories is currently lacking.

### CRISPR arrays are differently positioned and oriented for different Cas types

Most arrays tend to be close (< 10000 bp) to their respective Cas locus (Supplementary Fig. S2 and Table S4). However, there are huge differences between Cas types. Types I, II, and III tend to favor closeness with few exceptions, but arrays of type V and VI are almost always located farther away (> 500000 bp) from their Cas locus (Supplementary Fig. S7).

We analyze the position of CRISPR arrays in the genome with respect to their corresponding Cas locus by computing the distance between the closest array end and the respective Cas locus end. Thereby we determine if they are positioned “before” or “after” the Cas. Moreover, we determined the orientation of the arrays. We show both the distance and orientation (for arrays with close (< 10000 bp distance) Cas) for a subselection of interesting Cas types in Figure 2. Distributions for all types with sufficient data are shown in Supplementary Fig. S8.

**Fig. 2.**
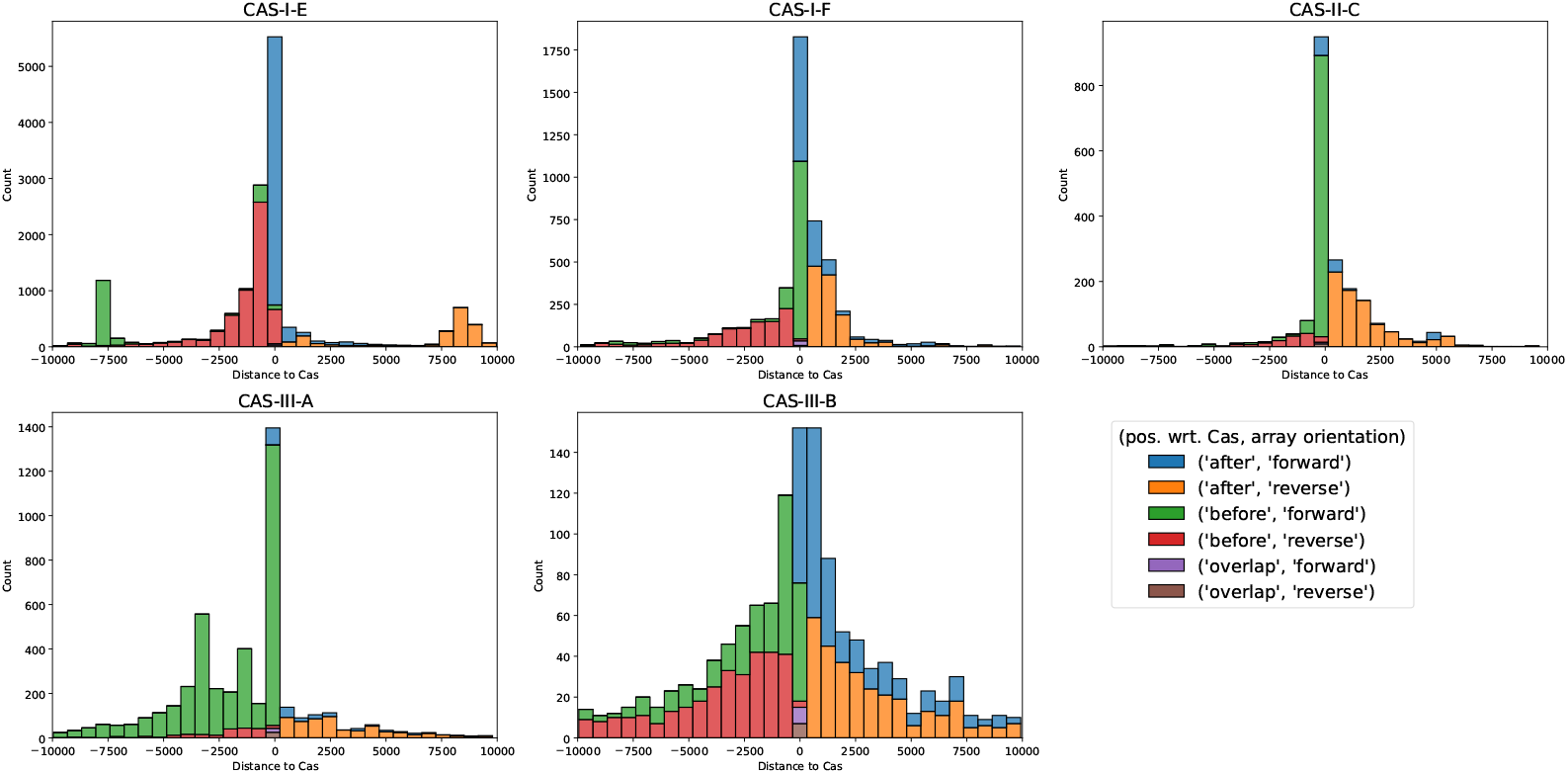
Distance to Cas and array orientation. We show the distance of CRISPR arrays to their respective Cas locus. The distances and labels ‘before’ and ‘after’ were obtained by comparing the positions of the arrays and Cas genes on the genome, i.e. arrays are positioned ‘before’/’after’ their Cas cassette and assigned negative/positive “Distance to Cas”, respectively. A small number of arrays are contained within their respective Cas locus and were labeled as ‘overlap’ (with distance 0). Additionally, we indicate the array orientation as predicted by CRISPRstrand. As expected, all types favor close CRISPR arrays, however the different Cas types show large differences in the distribution of distances and orientations with particular biases. In general, arrays close to the Cas locus tend to be forward oriented and more distantly located arrays tend to be reverse oriented. We identify multiple classes of distributions and show representative types for these behaviors here. Distributions for all Cas types with sufficient data can be found in Supplementary Fig. S8. In particular, type I-E has a bias for close arrays to be forward oriented and to be positioned after the Cas locus. Type I-E places close arrays on both sides of the Cas locus. I-E and II-A tend to favor to place the array before the Cas locus. However, II-A shows a significantly wider distribution. Lastly, type III-B shows significantly higher variance and places less importance of closeness to the Cas genes.

Closeness to the Cas locus tends to be favored for all types, and most arrays are less than 500 bp distant from a Cas locus. However, CRISPR types also show very different distributions of distances and distributions of orientations. Firstly, we found a tendency for all types that arrays close to Cas are forward oriented, whereas types further away from the Cas are reverse oriented. We grouped the different subtypes, with respect to their distance and orientation distributions, into multiple groups. For each group we show particularly interesting and representative types in Figure 2; marked **bold** in the text.

1. Type I-C, **I-E**, II-A, and IV-A tend to have arrays *after* and close to the Cas locus (in forward orientation), and arrays *before* the Cas tend to be more distant (in reverse orientation).
2. Type I-A, I-B, I-D, and **I-F** prefer close arrays *both before and after* the Cas locus (in forward orientation). More remote arrays are typically found *after* the Cas (in reverse orientation).
3. Type II-B, **II-C, III-A** tend to favor arrays arrays *before* and close to the Cas (in forward orientation). The more distant arrays are usually *after* the Cas (in reversed orientation). For III-A there are a substantial number of arrays more than 2500 bp away from the Cas locus.
4. Type **III-B** and III-D have a behavior similar to 2. However, they have distributions with much higher variance and less favoritism for specific orientations. This resembles a distribution one might expect in a setting with lower selection pressures that act on the orientation and the distance of CRISPR arrays and the associated Cas locus.
5. Although types V-A and VI-B are very abundant, very few of their arrays are close to their Cas. Where they are close, they show (similarly to 4) more random distributions (see Supplementary Fig. S8).

We further investigated the orientation and position of pairs of CRISPR arrays in genomes with only one Cas type and exactly two CRISPR arrays (Supplementary Fig. S9 and S10). Remarkably, we found that types I-D, I-E and I-F with two arrays almost exclusively position their arrays one on each side of the Cas locus. Possessing two arrays (or more) at the respective ends of the Cas locus is likely beneficial for various reasons. Multiple arrays provide redundancy, and their proximity to the Cas locus maintains high efficiency and enables joint regulation. Moreover, two arrays are likely to have more evenly distributed spacer expression compared to a single longer array, since spacers near their leader positions provide higher levels of immunity [8, 20].

Notably, I-E and I-F are very different in the distribution of array orientation. For I-E the pairs of arrays are almost exclusively both in forward or both in reverse orientation, whereas for type I-F the distribution of forward and reverse orientation of the two arrays seems to be independent of each other, regardless of whether the arrays are on the same or different sides of the Cas locus (Supplementary Fig. S10). I-D favors different orientations for pairs of arrays, but possesses a substantial number of array pairs with the same orientation (Supplementary Fig. S9). III-A behaves differently and typically places both arrays before the Cas and in the same (forward) orientation. The peaks around -2500 to -5000 in the distribution in Figure 2 for type III-A could thus indicate the beginning of a second array next to another array closer to the Cas locus.

Connecting the fact that I-E tends to have arrays in the same orientation with Figure 2, noting that most Cas loci of type I-E have one or two arrays, indicates that single arrays close to the Cas are most often after the Cas and forward oriented or before the Cas and reverse oriented. However, for array pairs one of the arrays is placed close before the Cas and reverse oriented or after the Cas and forward oriented and, quite frequently, the second array is much farther away and on the other side of the Cas (around 7500 bp) in the same orientation. In terms of colors in Figure 2: blue and green tend to form (forward oriented) pairs and red and orange tend to form (reverse oriented) pairs.

Our findings suggest that the distribution of array orientations are consistent within most types, with notable exceptions such as III-B. However, there is high variance in organisation between types, suggesting that the regulation of the transcription of Cas and arrays evolved to the specific needs of Cas types and host organisms [30].

Some systems, e.g. I-F, show heterogeneity in orientations even for arrays belonging to the same Cas locus (Supplementary Fig. S10 and [30]), which poses questions on how these systems with multiple closely located arrays (and their Cas) are regulated.

### Arrays vary in length across Cas types and are typically longer when located near Cas genes

The spacer array length tends to depend heavily on the Cas type, although most arrays have less than 50 spacers (average of 15.11, median of 9) and there is a small number of arrays with between 50 and 100 spacers (Supplementary Table S3 and Supplementary Fig. S11 and S12). In particular, I-A and I-D have remarkably large arrays with means of 51 and 42 spacers, respectively. Very large arrays are very rare, however, up to 246/317 spacers for type I-E/I-F and 587/587 for type I-B/I-U, respectively, exist. Types I (excluding I-A/I-D) and type III tend to have averages between 15 and 30, while type II has slightly smaller arrays with averages between 12 (II-C) and 18 (II-B) and types V and VI have far smaller arrays with averages and medians of 10 or lower. If the hypothesis that the loss rate of spacers is comparable between different types and species holds [2] this could indicate that type V and VI CRISPR systems are less often triggered to acquire new spacers, compared to type I, II, and III.

Splitting the set of all arrays regardless of the repeat sequence into two groups depending on how close they are to the Cas and redoing the statistics yields additional insight (Supplementary Table S4). We found that for most types, arrays close to their Cas are much longer than arrays further away. In particular, type II-C has a large difference between medians 12 (close) and 5 (not close). This holds for most types except I-A, III-B, III-D, V-F, VI-A and VI-C. Thus, one might be tempted to assume that distance to the Cas locus has a large impact on the spacer acquisition rate.

However, this distribution could also be the result of other effects. For example, leaderless arrays have been shown to tend to be shorter [18] and the predicted arrays that are further away could be more often leaderless. It could also be related to the absence of acquisition machinery (Cas1 and Cas2) in (close) Cas loci. In addition, when considering all CRISPR arrays within a genome, this will also include arrays with different repeats and misclassified or other CRISPR-like elements, which could cause the shift towards shorter arrays. In line with this argument, we find that arrays that are not close to a Cas locus tend to have diverse repeat sequences that differ from the repeats in arrays that are close (Supplementary Figure S15).

### Relative spacer acquisition rates for multiple arrays

We further investigated the insertion behavior by comparing the array lengths of arrays within the same genome. In contrast to the global array length distribution, when looking at the lengths of multiple arrays within the same genome, the deletion rate of spacers should be consistent across different arrays. This allows us to compare array lengths as a proxy for spacer acquisition rates within cells; see Methods and [2]. We call these comparative weights *insertion rate weights* and show their distribution in Figure 3. We restricted our analysis to genomes with exactly 2, 3 or 4 arrays to reduce other biases. For comparison, we computed neutral distributions of the insertion rate weights using parameters from the real dataset. In the neutral case, new spacers are inserted in any of the arrays with equal probability, which results in a multinomial distribution of the array lengths. See Methods for details about the derivation of the neutral distribution.

**Fig. 3.**
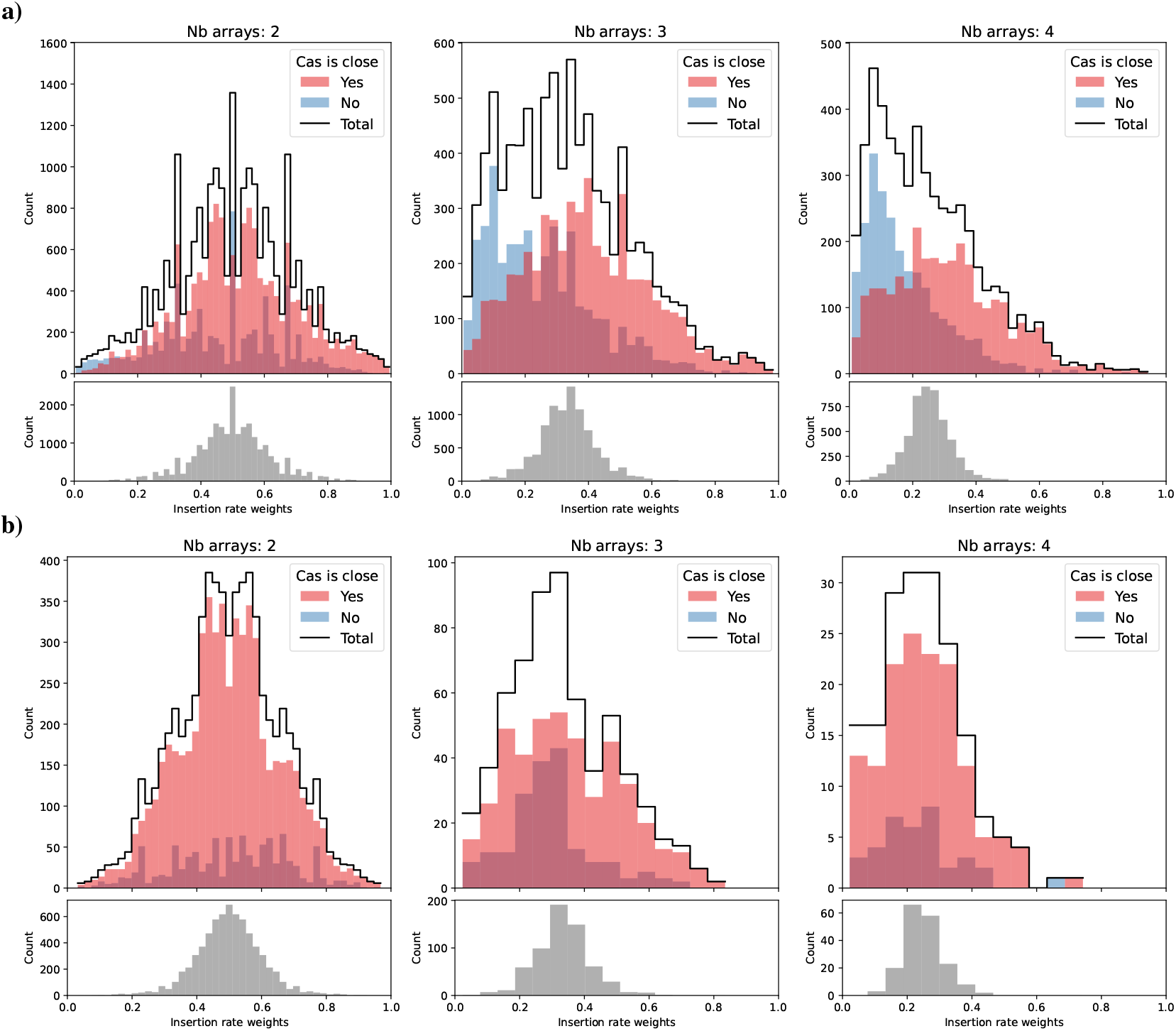
Distribution of the estimated insertion rate weights of arrays. At the top of each panel, we show the empirical count of insertion rate weights of arrays for fixed numbers of arrays (genomes with exactly 2, 3 or 4 arrays) colored red, if the respective Cas genes are nearer than 10000 bp, or colored blue otherwise (genomes with orphan arrays are excluded). The count plots are layered over each other (overlap in dark red). The total counts are shown in black. At the bottom of each panel, in gray, we show the expected neutral distribution of the insertion rate weights, if spacers are inserted at random in available arrays in the genome, for the given amount of samples. Details about the neutral distribution are described in Methods. Note, that the counts (y-axis) are scaled differently for the observed and expected distribution. **a)** shows the weights across the whole dataset and **b)** only for arrays with the same consensus repeat. Note that in a), the distribution of arrays which are further away (dark and light red) dominate the the left side of the plots and the distribution of arrays closer to their Cas genes (blue and dark red) are further to the right. This indidates that arrays close to their Cas genes are generally larger than arrays further away. In b), most arrays are close to the Cas genes, but there is no bias of arrays away from the Cas locus to be longer. This indicates that, even though distance from the Cas genes is relevant for the acquisition rate of arrays, possessing the same consensus repeat is of even greater importance.

As expected, in the neutral case, when all arrays have a similar chance to acquire new spacers, and thus similar lengths, the relative weight concentrates around the inverse of the number of arrays. However, in Figure 3a, we see that the overall insertion rate weights are distributed around the expected 0.5/0.33 for 2/3 arrays, respectively, but not around 0.25 for 4 arrays. The distribution has an increasing weight towards the lower end indicating fewer large and more small arrays. In general, the observed distributions have significantly higher variance and are wider. For the arrays close to their Cas (light red/dark red) and the arrays far away from their Cas (blue/dark red) we noted that the distribution of relative lengths is shifted and arrays with close Cas tend to have larger weights than those with a more distant Cas locus. This can also be seen in the distribution of array lengths with respect to their proximity to Cas (Supplementary Table S4, Supplementary Fig. S11 and S12).

One might be tempted to conclude from this pattern that spacer acquisition is more efficient if the acquisition machinery is positioned closer to the array. However, when we further reduced the available arrays to arrays with the same consensus repeat we found that the left shift of the distributions vanishes. In Figure 3b, the distribution of arrays more distant to the Cas locus is entirely enveloped by the distribution for close to Cas arrays. Moreover, more weight of the distribution is around 0.5/0.33/0.25 as expected for equally distributed acquisitions among the 2/3/4 arrays and the variance is closer to the neutral expectation.

This effect appears to be Cas gene-specific, as shown in Supplemental Table S4: some types exhibit a large decrease in (average) array length for distant arrays, while others show no noticeable decrease.

Nonetheless, this indicates that although the distance to the Cas is correlated with array length, possessing the same (consensus) repeat is far more important for consistent acquisition of new spacers. Consequently, arrays with the same repeat often maintain comparable lengths and are equally likely updated regardless of their distance to the Cas locus.

The relevance of the (first) repeat for spacer acquisition is supported by recent findings that Cas1-Cas2 complexes show higher insertion efficiency by matching to parts of the leader and the first repeat for acquisition for type II-A [31].

Moreover, repeats were found to be relevant for the processing of crRNA in type II systems. It was found that the leader forms a stem-loop structure with the first leader adjacent repeat which increases crRNA processing efficiency at the leader end and inhibits creation of non-functional ‘extraneous’ crRNA [17, 32]. Note that we only analyze the acquisition rates for new spacers, and we can not infer the expression rates of these spacers as crRNA and thus draw direct conclusions about the inferred immunity or the impact of Cas distance for expression efficiency.

Lastly, possessing the same consensus repeat could also be a proxy measure for the arrays to have the same leaders, which have been found to be crucial for acquisition and expression of CRISPR arrays.

We are aware of only one experimental setup in which acquisitions split among two arrays are considered. While the authors also found that both arrays are used for acquisition, the relative acquisition rates are hard to disentangle from a potential copy number bias towards the origin of replication in a replicating bacterial culture [33].

### Repeat distances cluster according to Cas types

We computed the (normalized) Levenshtein distance between consensus repeats of the dataset. We found that, in general, the distribution of the normalized consensus repeat distances is bimodal (or even multimodal) with one focal point around zero and one or even more focal points in the distance range 0.4 - 0.6 for increasing number of arrays and types (Supplementary Fig. S13). Multiple arrays close to the same Cas locus frequently occur, and in this case it is likely beneficial to maintain similar repeats. Moreover, some types are known to use the acquisition machinery of other Cas loci and thus might benefit from similar repeats. Both cases result in the mode around 0. In general, repeat similarity might not be necessary or even detrimental when supporting multiple CRISPR-Cas systems with different tasks. This and an increasing number of co-occurring different types of Cas for which only a limited number are compatible likely leads to the additional modes. To characterize and compare the repeats within Cas types and for different Cas types further, we performed MDS, with respect to the pairwise distances of the consensus repeats, to project the repeats of each array into a 2-d space; see Figure 4 and Supplementary Fig. S14, S15.

**Fig. 4.**
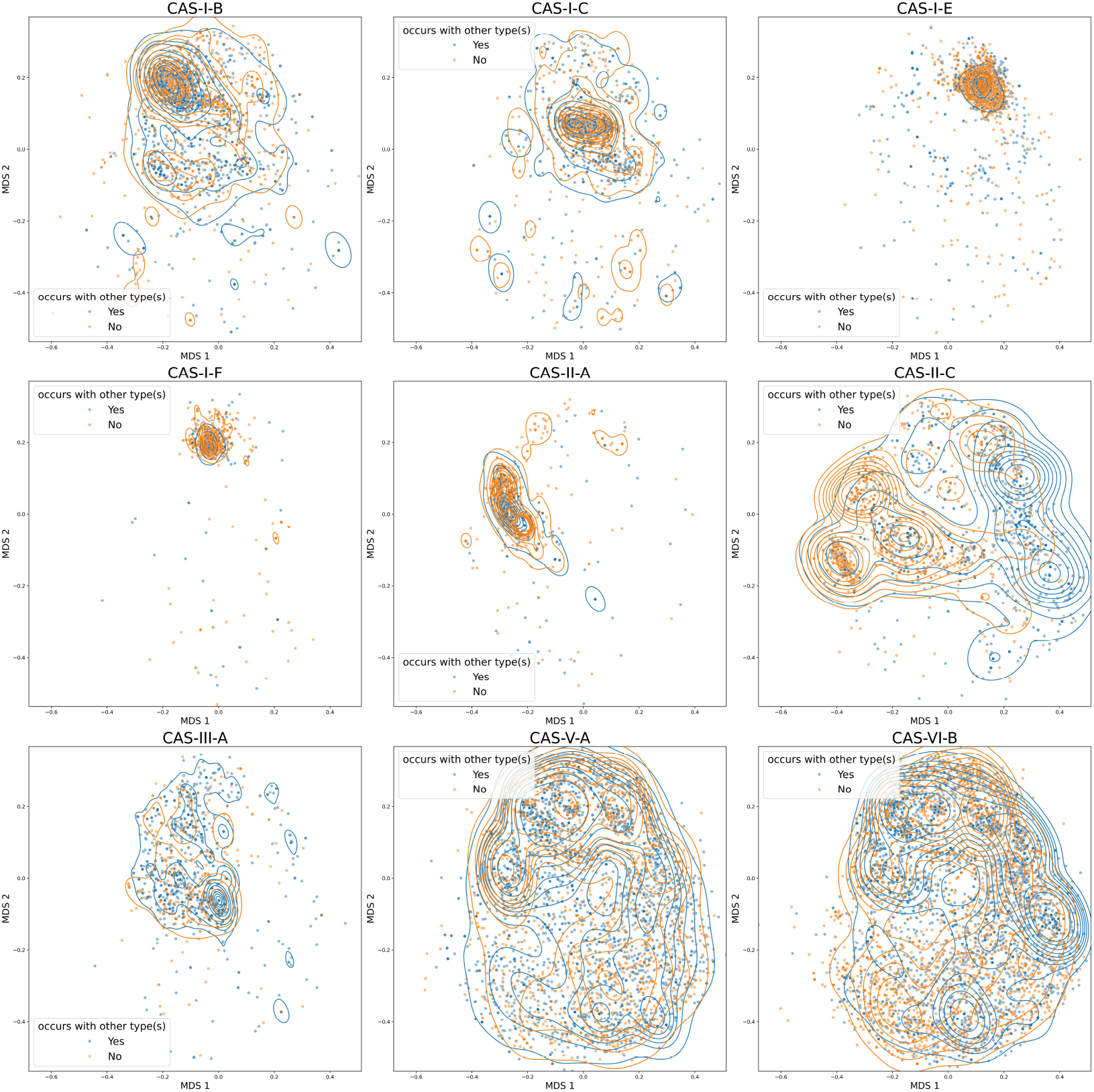
Visualization of repeat similarity by projection into 2-d space with multidimensional scaling. The plot shows the projection of the consensus repeats (for each array) into 2-d space with multi dimensional scaling with respect to the pairwise consensus repeat edit distances (see Methods for details). The MDS is done globally across all arrays and types, i.e. all shown figures are using the same MDS projection and thus are in the same coordinate system. However, we separated the Cas types into subplots for visual clarity. Each dot corresponds to the consensus repeat of one array and is colored blue, if other Cas types were found within the same genome, and orange otherwise. Each single dot is semi-transparent, thus more intensive color indicates multiple arrays with the same consensus repeat. To illustrate where the distributions are most concentrated, we show the contours of kernel density estimates for both co- and non-co-occurring types in their respective colors. We show only a selection of representative Cas types here; see Supplementary Fig. S14 for plots of all Cas types. The types show large differences in their distributions. In particular, type I arrays tend to have very concentrated distinguishable distributions and have closely related repeats. However, type II shows much more variance, in particular, type II-C shows multiple distinguishable clusters. Type V-A and VI-B are distributed across the whole space and seem to share similar repeats with all other types.

MDS is a method to translate the pairwise distances of a set of points (here the consensus repeats of each array) into a less-dimensional (here 2-d) space such that the pairwise distances are maintained as well as possible. Thus, MDS allows us to visualize the similarity of the consensus repeats losing as little information about their edit distance as possible; see Methods for details about the MDS. Note that the MDS was performed using all arrays, and all Figures are created using the same MDS projection, only different subsets of arrays are shown for visual clarity.

Most type I repeats form clearly distinguishable clusters with some outliers (which occur for all types) that are possibly misclassified arrays. In Supplementary Fig. S15 which shows the repeat MDS colored according to the distance from their Cas locus, it becomes clear that many repeat sequence outliers are also far away from their Cas locus. This further reinforces that these arrays might be misclassified as the wrong type, inactive, or false CRISPR detections [34, 35].

However, it is remarkable that types I-B, I-E and I-F, although they have more similar repeats, co-occur only rarely (Figure 1) with I-E and I-F in particular avoiding most other Cas types, excluding some of the widely spread V and VI subtypes. Type V and VI show very wide distributions that populate almost the whole MDS space which is very similar to the seemingly random distribution exhibited by the orphan arrays (Supplementary Fig. S14). This could suggest that they are very widely compatible with other Cas types, an idea that is supported by their relative frequent co-occurence with other types seen in Figure 1 and experimental studies [13, 36] who found that a VI-B array used the acquisition machinery of a II-C locus. However, it can also suggest that many type V and VI arrays might be misclassified or false CRISPR detections [34, 35].

Types II tend to fall somewhere in between types I and V/VI showing wider distributions with some still distinguishable clusters. In particular, type II-C shows multiple clusters.

### Type II-C repeats cluster into subgroups

In Figure 5, we show a more detailed breakdown of the MDS projections with marginal distributions for a subgroup of types. For II-C in Figure 5a there are clear differences between repeats of II-C that predominantly co-occur with other types and smaller clusters where no other Cas type is present. This is consistent with the classification by Makarova et al. [4] where they found two subtypes of II-C arrays. However, considering Supplementary Fig. S15, it becomes clear that arrays in the cluster on the right, which co-occurs with other Cas types, are mostly far away from their Cas locus and thus have remarkably small arrays with often less than 5 spacers, see Supplementary Table S4 and Supplementary Fig. S12. It is possible that these are misclassifications, however, none of the other Cas types shows similar behavior, and their close clustering suggests that this might be a special case for type II-C.

**Fig. 5.**
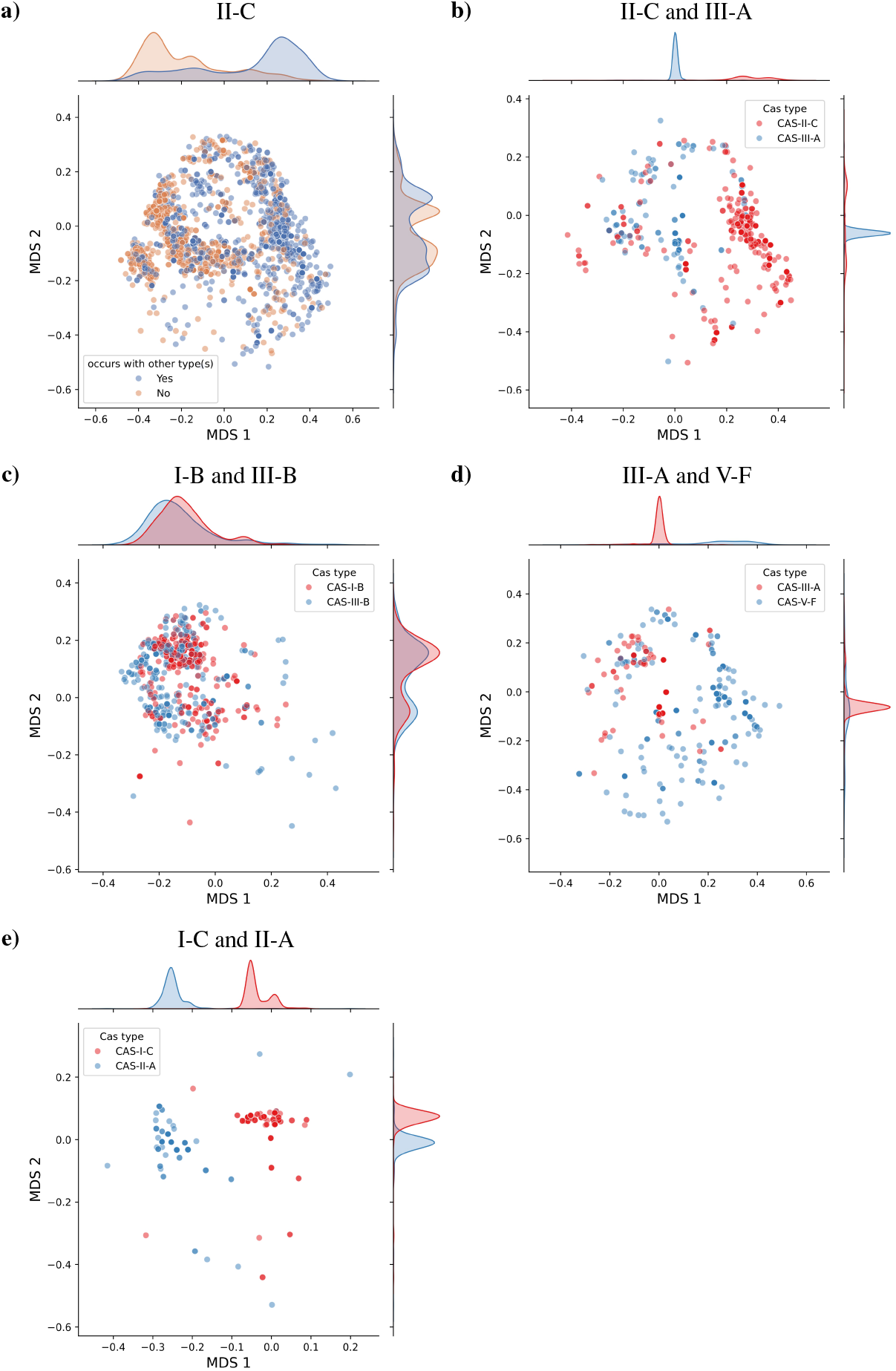
Repeat similarity MDS joint scatterplot and marginal distributions. In this plot, we show the MDS projections of the repeats shown in Figure 4 with additional kernel density estimates of the marginal distributions for both MDS coordinates for interesting subsets of arrays. See methods for details about the MDS and kernel density estimates. **a)** shows only Cas-II-C which clearly shows the existence of different subgroups of repeats. Moreover, we see that they can be separated into groups which occur more frequently and less frequently with other Cas types. **b) – e)** show only the repeats of arrays in genomes that posess arrays of both Cas types that have been found to significantly co-occur (see Figure 1), as noted in each title. **b)** shows that co-occurring II-C and III-B repeats are most often distant from each other. Most II-C repeats are found in the right cluster and are mostly far away from their Cas and shorter in length (see Supplementary Fig. S12, S15). In **c)**, repeats of types I-B and III-B are very similar and similarly distributed with almost complete overlap. In **d)**, some arrays of type III-A and V-F have similar repeats, but in general are relatively distant from each other, and V-F has a much wider distribution than III-A. I-C and II-A in **e)** clearly have distant distinct repeats.

The type II-C system has been shown to be peculiar in multiple ways, e.g. they acquire spacers at the 3’ end [13, 36–38], carry promoters in the repeats [13, 37, 39] and show unique characteristics in crRNA expression [32]. Moreover, to date, orientation tools fail to reach a consensus on the orientation of many type II-C systems [40]. These particularities point towards the unique functionality of type II-C systems, potentially resulting in multiple distinct subtypes and the pattern observed here.

### Repeat similarity of co-occurring CRISPR systems

In Figure 5b–e, we show the repeat MDS projections for pairs of CRISPR types with high co-occurrence rate (Figure 1). They show a differentiated picture, suggesting that beneficial co-occurrence of types can be efficient for very varied repeat agreement.

We see in Figure 5b that type III-A has a high overlap with the middle II-C cluster. However, type II-C arrays with cooccuring types (and in particular III-A) are mostly found within the right cluster. Thus, II-C and III-A repeats that co-occur have, in most cases, different repeats.

In general, repeat similarity could be important for efficient spacer acquisition in trans. I-B and III-B clearly have very similar almost exactly overlapping distributions for their repeats (Figure 5c). Makarova et al. [4] found that I-B has spacer acquisition genes, like Cas1 and Cas2, and III-B does not. This suggests that III-B uses the acquisition machinery of I-B. Vink et al. found results consistent with this in the distribution of spacers and targets of arrays of both types [9]. Similarly, plasmid-encoded adaptation-deficient type IV systems have also been found to frequently co-occurr with host-encoded adaptation-competent systems with leader, repeat and PAM similarity [26].

III-A and V-F share similar repeats for only a subset of cooccurring arrays (Figure 5d). According to Makarova et al. [4] both type III-A and some subtypes of V-F do not necessarily possess integral genes for spacer integration such as Cas1 and Cas2. This suggests that it might be beneficial for III-A and V-F to maintain similar repeats and pool their Cas machinery to allow for new spacer acquisition and to retain functionality. It is remarkable that, in both Figures 5b and d, III-A is shown to have a very concentrated distribution of repeats, but co-occurs with much more universalist types in terms of repeat diversity and negative aloneness score (II-C, V-F, VI-B, Figure 1). Moreover, Hoikkala et al. [13] show that for II-C and VI-B – arrays with only limited repeat similarity – VI-B can acquire spacers using the II-C machinery.

I-C and II-A are remarkable in the sense that their repeats are related, but they do not overlap in Figure 5e. Moreover, they are the types with the highest co-occurrence ratio and according to Makarova et al. [4] both have functional genes for spacer acquisition. This suggests that these types fulfill complementary and beneficial functions but operate independently for spacer acquisition.

CRISPR systems could also cooperate during interference, utilizing other Cas genes in trans or employing spacers or crRNA from other arrays in their effector complexes. The co-opting of a single Cas6 protein to process crRNA for both a type I-A and two III-B systems was experimentally observed in *Sulfolobus islandicus* [41]. There is also extensive experimental evidence for the sharing of crRNA between DNA and RNA targeting systems which could provide sustained defense against phages that employ anti-CRISPR tactics [42]. Majumdar et al. found crRNA from all 7 arrays in the genome of *Pyrococcus furiosus* in the effector complexes of the 3 co-occurring Cas loci of types I-A, I-G and III-B [42].

Additionally, different CRISPR systems could serve different roles at different stages of phage defense. For example, Silas et al. found that the type III-B system in *Marinomonas mediterranea* uses crRNA from type I-F arrays in the same system [12]. They propose that the III-B system serves as a backup for the I-F system in the event of phage evasion via PAM mutation since the III-B system is not PAM dependent. Moreover, the type VI interference mechanism (Cas13) induces collateral RNA degradation when a certain amount of target RNA is detected within the cell, rapidly resulting in cell dormancy or death [43]. Thus, the frequently co-occurring VI-B systems could serve as a “fall-back” system for the cell in the event of catastrophic infection with an invasive element when other more specific CRISPR systems have failed to stop the infection.

### Summary and open questions

In Table 1 we give a short summary of our findings, relevant subtypes where these findings are observable, and state some of the possible mechanisms underlying the observations as suggested in the literature.

**Table 1.**
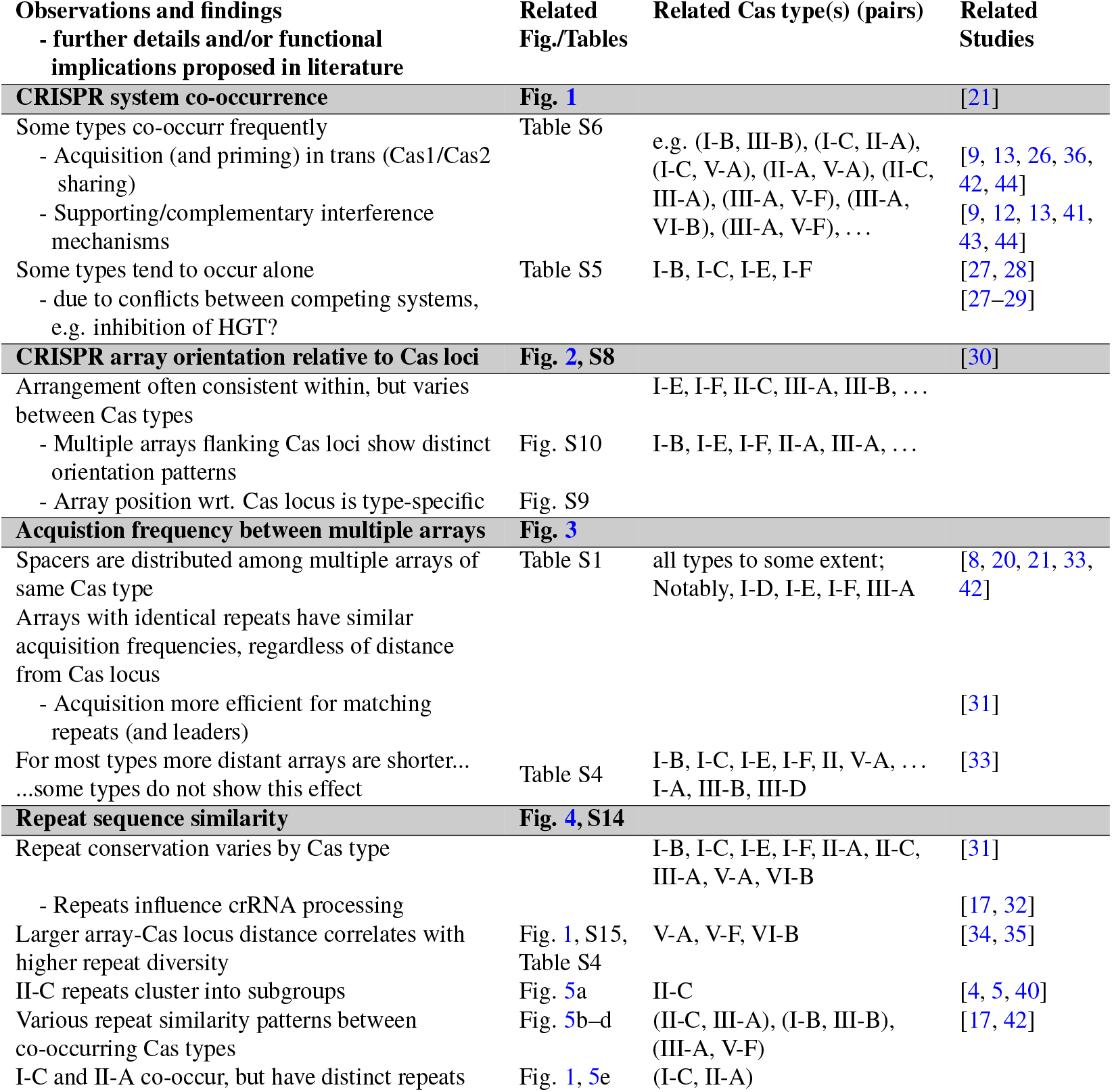
Summary of findings and possible functional implications proposed/supported in the literature. Here we summarize our main findings in this mansucript. The findings are separated into 4 sections marked in bold. For each finding, we provide the relevant figures along with related studies and literature references for the *entire* section. Within these sections, the findings are listed in greater detail. Indented we give some additional details and/or functional implications that arise from the observations and are proposed and/or supported by the literature shown in related studies. For each point we give additional relevant figures and tables and state relevant Cas type(s) (pairs) for which the findings are most evident.

The most notable fact is the sheer complexity of the dynamics of CRISPR system co-occurrence and interaction, additionally to the large diversity of CRISPR types in their array and Cas configuration. Naturally, interaction between systems are hard to explore in empirical studies since they introduce additional complexity and biases. However, it is important to understand why some systems occur more frequently together (Figure 1), what the benefit of the co-occurrence of systems is, and through which mechanisms these systems can interact.

Figure 1 and Supplementary Table S5 offer candidates for such interacting systems that could merit detailed experimental investigation. Moreover, Figure 5 illustrates that these systems might interact on different levels of acquisition, expression, and interference.

We mainly focused on genomes with a limited number of Cas types and arrays to mitigate the impact of statistical noise. However, the large number of genomes with diverse Cas systems require further investigation to better understand their interaction. Recent studies such as [13, 44] and our results indicate that the interaction between systems is crucial and underexplored. Margolis and Meeske investigated the interactions between three CRISPR systems of type I-B, II-C and VI in *Listeria seeligri*. They observed that type VI uses II-C adaptation to insert spacers and I-B primes the acquisitions of VI and II-C [44]. This suggests that communication between multiple systems might be more complex than previously expected.

There are many other open questions that go beyond our findings, such as the impact of the distance to the Cas locus on expression and interference efficiency, as well as the impact of Cas and array orientation on acquisition, expression, and interference efficiency.

Together with these results, our findings, including the observation that arrays with identical repeat have similar acquisition activity across vast genomic distances, suggest that we require a more holistic approach considering all CRISPR systems across the genome to understand their function and interaction. Furthermore, the purpose of possessing multiple (for some types very close) CRISPR arrays with various orientation configurations are uncertain. Although several hypotheses have been proposed to explain the distribution of spacers across multiple arrays [8, 21], much remains to be understood — particularly through experimental studies — about how spacer acquisition, deletion, and interference proceed in systems harboring multiple arrays and Cas types.

## Conclusion

We show that the arrays of CRISPR-Cas systems have a remarkable diversity that can diverge wildly between types. However, some clear patterns emerge. Although array length decreases for arrays distant from the next Cas locus, hinting at a low spacer acquisition rate, this effect almost completely vanishes if the repeats of the arrays agree. This suggests that arrays with identical repeats are equally likely to acquire new spacers, regardless of their genomic position and distance to the Cas locus. Moreover, types show clear distinct patterns of positioning and orienting their arrays around the Cas locus. The observed variations might be a result of the selective pressure to diversify the bacterial defense reportoire to avoid or compensate anti-CRISPR mechanisms and persist in the evolutionary arms race with phages.

The different patterns in the frequency of co-occurrence of Cas types and the differences in repeat (dis)similarity and array orientation identified here invite further exploration. Understanding the underlying mechanisms shaping these patterns – including spacer acquisition and inference in trans, layered defense strategies, and redundancies within prokaryotic immune systems – will help to assess the synergies and incompatibilities of prokaryotic defense systems.

## ACKNOWLEDGEMENTS

AF and FB are funded by the Deutsche Forschungsgemeinschaft (DFG, German Research Foundation) under the Priority Program - SPP 2141 - Project number Ba-5529/1-1. FB is funded by the Deutsche Forschungsgemeinschaft (DFG, German Research Foundation) under Germany’s Excellence Strategy – EXC number 2064/1 – Project number 390727645, and EXC 2124 – Project number 390838134.

This work was supported by German Research Foundation (DFG) [BA 2168/23-1 and BA 2168/23-2 SPP 2141 ]; Much more than Defence: the Multiple Functions and Facets of CRISPR–Cas; Baden-Wuerttemberg Ministry of Science, Research and Art; University of Freiburg.

This work was supported by the BMBF-funded de.NBI Cloud within the German Network for Bioinformatics Infrastructure (de.NBI) (031A532B, 031A533A, 031A533B, 031A534A, 031A535A, 031A537A, 031A537B, 031A537C, 031A537D, 031A538A).

## Notes

### Competing Interest Statement

The authors have declared no competing interest.

### Summary of Updates

Added discussion and additional references where suitable; added additional technical details; adjusted Figures to be more readable (Figure 4 in particular); clarified and improved writing; corrected typos; added some supplemental figures and improved readibility; updated author affiliations.

## Bibliography

1. Edze R. Westra, Andrea J. Dowling, Jenny M. Broniewski, and Stineke van Houte. Evolution and Ecology of CRISPR. Annual Review of Ecology, Evolution, and Systematics, 47(1): 307–331, 2016. doi: 10.1146/annurev-ecolsys-121415-032428.

2. Axel Fehrenbach, Alexander Mitrofanov, Omer S Alkhnbashi, Rolf Backofen, and Franz Baumdicker. SpacerPlacer: ancestral reconstruction of CRISPR arrays reveals the evolutionary dynamics of spacer deletions. Nucleic Acids Research, gkae772, 2024. doi: 10.1093/nar/gkae772.

3. Adrián López-Beltrán, João Botelho, and Jaime Iranzo. Dynamics of CRISPR-mediated virus-host interactions in the human gut microbiome. bioRxiv, 10.1101/2024.01.23.576851, 2024. doi: 10.1101/2024.01.23.576851.

4. Kira S. Makarova, Yuri I. Wolf, Jaime Iranzo, Sergey A. Shmakov, Omer S. Alkhnbashi, Stan J. J. Brouns, Emmanuelle Charpentier, David Cheng, Daniel H. Haft, Philippe Horvath, Sylvain Moineau, Francisco J. M. Mojica, David Scott, Shiraz A. Shah, Virginijus Siksnys, Michael P. Terns, Česlovas Venclovas, Malcolm F. White, Alexander F. Yakunin, Winston Yan, Feng Zhang, Roger A. Garrett, Rolf Backofen, John Van Der Oost, Rodolphe Barrangou, and Eugene V. Koonin. Evolutionary classification of CRISPR–Cas systems: a burst of class 2 and derived variants. Nature Reviews Microbiology, 18(2):67–83, 2020. doi: 10.1038/s41579-019-0299-x.

5. Kira S. Makarova, Sergey A. Shmakov, Yuri I. Wolf, Pascal Mutz, Han Altae-Tran, Chase L. Beisel, Stan J. J. Brouns, Emmanuelle Charpentier, David Cheng, Jennifer Doudna, Daniel H. Haft, Philippe Horvath, Sylvain Moineau, Francisco J. M. Mojica, Patrick Pausch, Rafael Pinilla-Redondo, Shiraz A. Shah, Virginijus Siksnys, Michael P. Terns, Jesse Tordoff, Česlovas Venclovas, Malcolm F. White, Alexander F. Yakunin, Feng Zhang, Roger A. Garrett, Rolf Backofen, John Van Der Oost, Rodolphe Barrangou, and Eugene V. Koonin. An updated evolutionary classification of CRISPR–Cas systems including rare variants. Nature Microbiology, 2025. doi: 10.1038/s41564-025-02180-8.

6. Rafael Pinilla-Redondo, Jakob Russel, David Mayo-Muñoz, Shiraz A Shah, Roger A Garrett, Joseph Nesme, Jonas S Madsen, Peter C Fineran, and Søren J Sørensen. CRISPR-Cas systems are widespread accessory elements across bacterial and archaeal plasmids. Nucleic Acids Research, 50(8):4315–4328, 2022. doi: 10.1093/nar/gkab859.

7. Michael Zaayman and Rachel M. Wheatley. Fitness costs of CRISPR-Cas systems in bacteria. Microbiology, 168(7), 2022. doi: 10.1099/mic.0.001209.

8. Jl. Weissman, William F. Fagan, and Philip L.F. Johnson. Selective Maintenance of Multiple CRISPR Arrays Across Prokaryotes. The CRISPR Journal, 1(6):405–413, 2018. doi: 10.1089/crispr.2018.0034.

9. Jochem N. A. Vink, Jan H. L. Baijens, and Stan J. J. Brouns. PAM-repeat associations and spacer selection preferences in single and co-occurring CRISPR-Cas systems. Genome Biology, 22(1):281, 2021. doi: 10.1186/s13059-021-02495-9.

10. Susanne Erdmann and Roger A. Garrett. Selective and hyperactive uptake of foreign DNA by adaptive immune systems of an archaeon via two distinct mechanisms. Molecular Microbiology, 85(6):1044–1056, 2012. doi: 10.1111/j.1365-2958.2012.08171.x.

11. Lisa Nickel, Weidenbach, Katrin, Jäger, Dominik, Backofen, Rolf, Lange Sita J., Heidrich, Nadja,, and Ruth A. Schmitz. Two CRISPR-Cas systems inMethanosarcina mazeistrain Gö1 display common processing features despite belonging to different types I and III. RNA Biology, 10(5):779–791, 2013. doi: 10.4161/rna.23928.

12. Sukrit Silas, Patricia Lucas-Elio, Simon A. Jackson, Alejandra Aroca-Crevillén, Loren L. Hansen, Peter C. Fineran, Andrew Z. Fire, and Antonio Sánchez-Amat. Type III CRISPR-Cas systems can provide redundancy to counteract viral escape from type I systems. eLife, 6: e27601, 2017. doi: 10.7554/eLife.27601.

13. Ville Hoikkala, Janne Ravantti, César Díez-Villaseñor, Marja Tiirola, Rachel A. Conrad, Mark J. McBride, Sylvain Moineau, and Lotta-Riina Sundberg. Cooperation between Different CRISPR-Cas Types Enables Adaptation in an RNA-Targeting System. mBio, 12(2):e03338– 20, 2021. doi: 10.1128/mBio.03338-20.

14. Shayna R Deecker and Alexander W Ensminger. Type I-F CRISPR-Cas Distribution and Array Dynamics in Legionella pneumophila. G3 Genes|Genomes|Genetics, 10(3):1039–1050, 2020. doi: 10.1534/g3.119.400813.

15. Chhandosee Ganguly, Saadi Rostami, Kole Long, Swarmistha Devi Aribam, and Rakhi Rajan. Unity among the diverse RNA-guided CRISPR-Cas interference mechanisms. Journal of Biological Chemistry, 300(6):107295, 2024. doi: 10.1016/j.jbc.2024.107295.

16. Jon McGinn and Luciano A. Marraffini. CRISPR-Cas Systems Optimize Their Immune Response by Specifying the Site of Spacer Integration. Molecular Cell, 64(3):616–623, 2016. doi: 10.1016/j.molcel.2016.08.038.

17. Chunyu Liao, Sahil Sharma, Sarah L. Svensson, Anuja Kibe, Zasha Weinberg, Omer S. Alkhnbashi, Thorsten Bischler, Rolf Backofen, Neva Caliskan, Cynthia M. Sharma, and Chase L. Beisel. Spacer prioritization in CRISPR-Cas9 immunity is enabled by the leader RNA. Nature Microbiology, 7(4):530–541, 2022. doi: 10.1038/s41564-022-01074-3.

18. Omer S Alkhnbashi, Shiraz A Shah, Roger A Garrett, Sita J Saunders, Fabrizio Costa, and Rolf Backofen. Characterizing leader sequences of CRISPR loci. Bioinformatics, 32(17): i576–i585, 2016. doi: 10.1093/bioinformatics/btw454.

19. Murat Buyukyoruk, Pushya Krishna, Andrew Santiago-Frangos, and Blake Wiedenheft. Discovery of Diverse CRISPR Leader Motifs, Putative Functions, and Applications for Enhanced CRISPR Detection and Subtype Annotation. The CRISPR Journal, crispr.2024.0093, 2025. doi: 10.1089/crispr.2024.0093.

20. Alexander Martynov, Konstantin Severinov, and Iaroslav Ispolatov. Optimal number of spacers in CRISPR arrays. PLOS Computational Biology, 13(12):e1005891, 2017. doi: 10.1371/journal.pcbi.1005891.

21. Aude Bernheim, David Bikard, Marie Touchon, and Eduardo PC Rocha. Atypical organizations and epistatic interactions of CRISPRs and cas clusters in genomes and their mobile genetic elements. Nucleic Acids Research, gkz1091, 2019. doi: 10.1093/nar/gkz1091.

22. Alexander Mitrofanov, Omer S Alkhnbashi, Sergey A Shmakov, Kira S Makarova, Eugene V Koonin, and Rolf Backofen. CRISPRidentify: identification of CRISPR arrays using machine learning approach. Nucleic Acids Research, 49(4):e20–e20, 2021. doi: 10.1093/nar/gkaa1158.

23. Victor A Padilha, Omer S Alkhnbashi, Shiraz A Shah, André CPLF de Carvalho, and Rolf Backofen. CRISPRcasIdentifier: Machine learning for accurate identification and classification of CRISPR-Cas systems. GigaScience, 9(6):giaa062, 2020. doi: 10.1093/gigascience/giaa062.

24. Omer S. Alkhnbashi, Fabrizio Costa, Shiraz A. Shah, Roger A. Garrett, Sita J. Saunders, and Rolf Backofen. CRISPRstrand: predicting repeat orientations to determine the crRNA-encoding strand at CRISPR loci. Bioinformatics, 30(17):i489–i496, 2014. doi: 10.1093/bioinformatics/btu459.

25. Yi Wu, Sofya K. Garushyants, Anne van den Hurk, Cristian Aparicio-Maldonado, Simran Krishnakant Kushwaha, Claire M. King, Yaqing Ou, Thomas C. Todeschini, Martha R. J. Clokie, Andrew D. Millard, Yilmaz Emre Gençay, Eugene V. Koonin, and Franklin L. Nobrega. Bacterial defense systems exhibit synergistic anti-phage activity. Cell Host & Microbe, 32(4):557–572.e6, 2024. doi: 10.1016/j.chom.2024.01.015.

26. Rafael Pinilla-Redondo, David Mayo-Muñoz, Jakob Russel, Roger A Garrett, Lennart Randau, Søren J Sørensen, and Shiraz A Shah. Type IV CRISPR–Cas systems are highly diverse and involved in competition between plasmids. Nucleic Acids Research, 48(4):2000–2012, 2020. doi: 10.1093/nar/gkz1197.

27. Marie Touchon and Eduardo P. C. Rocha. The Small, Slow and Specialized CRISPR and Anti-CRISPR of Escherichia and Salmonella. PLoS ONE, 5(6):e11126, 2010. doi: 10.1371/journal.pone.0011126.

28. Cristóbal Almendros, Noemí M. Guzmán, Jesús García-Martínez, and Francisco J. M. Mojica. Anti-cas spacers in orphan CRISPR4 arrays prevent uptake of active CRISPR–Cas I-F systems. Nature Microbiology, 1(8):16081, 2016. doi: 10.1038/nmicrobiol.2016.81.

29. Yang Liu, João Botelho, and Jaime Iranzo. Timescale and genetic linkage explain the variable impact of defense systems on horizontal gene transfer. Genome Research, genome;gr.279300.124v2, 2025. doi: 10.1101/gr.279300.124.

30. Ognjen Milicevic, Jelena Repac, Bojan Bozic, Magdalena Djordjevic, and Marko Djordjevic. A Simple Criterion for Inferring CRISPR Array Direction. Frontiers in Microbiology, 10:2054, 2019. doi: 10.3389/fmicb.2019.02054.

31. Jenny G Kim, Sandra Garrett, Yunzhou Wei, Brenton R Graveley, and Michael P Terns. CRISPR DNA elements controlling site-specific spacer integration and proper repeat length by a Type II CRISPR–Cas system. Nucleic Acids Research, 47(16):8632–8648, 2019. doi: 10.1093/nar/gkz677.

32. Maximilian Feussner, Angela Migur, Alexander Mitrofanov, Omer S Alkhnbashi, Rolf Backofen, Chase L Beisel, and Zasha Weinberg. Disparate mechanisms counteract extraneous CRISPR RNA production in type II-C CRISPR-Cas systems. microLife, 6:uqaf007, 2025. doi: 10.1093/femsml/uqaf007.

33. Luke J. Peach, Haoyun Zhang, Brian P. Weaver, and James Q. Boedicker. Assessing spacer acquisition rates in E. coli type I-E CRISPR arrays. Frontiers in Microbiology, 15, 2025. doi: 10.3389/fmicb.2024.1498959.

34. Quan Zhang and Yuzhen Ye. Not all predicted CRISPR–Cas systems are equal: isolated cas genes and classes of CRISPR like elements. BMC Bioinformatics, 18(1):92, 2017. doi: 10.1186/s12859-017-1512-4.

35. Murat Buyukyoruk, William S. Henriques, and Blake Wiedenheft. Clarifying CRISPR: Why Repeats Identified in the Human Genome Should Not Be Considered CRISPRs. The CRISPR Journal, 6(3):216–221, 2023. doi: 10.1089/crispr.2022.0106.

36. Elina Laanto, Ville Hoikkala, Janne Ravantti, and Lotta-Riina Sundberg. Long-term genomic coevolution of host-parasite interaction in the natural environment. Nature Communications, 8(1):111, 2017. doi: 10.1038/s41467-017-00158-7.

37. Yan Zhang, Nadja Heidrich, Biju Joseph Ampattu, Carl W. Gunderson, H. Steven Seifert, Christoph Schoen, Jörg Vogel, and Erik J. Sontheimer. Processing-Independent CRISPR RNAs Limit Natural Transformation in Neisseria meningitidis. Molecular Cell, 50(4):488–503, 2013. doi: 10.1016/j.molcel.2013.05.001.

38. Alexander Mitrofanov, Chase L. Beisel, Franz Baumdicker, Omer S. Alkhnbashi, and Rolf Backofen. Comprehensive Analysis of CRISPR Array Repeat Mutations Reveals Subtype-Specific Patterns and Links to Spacer Dynamics. bioRxiv, 2025. doi: 10.1101/2025.04.02.646798.

39. Gaurav Dugar, Ryan T. Leenay, Sara K. Eisenbart, Thorsten Bischler, Belinda U. Aul, Chase L. Beisel, and Cynthia M. Sharma. CRISPR RNA-Dependent Binding and Cleavage of Endogenous RNAs by the Campylobacter jejuni Cas9. Molecular Cell, 69(5):893–905.e7, 2018. doi: 10.1016/j.molcel.2018.01.032.

40. Axel Fehrenbach, Alexander Mitrofanov, Omer S. Alkhnbashi, Rolf Backofen, and Franz Baumdicker. An evolutionary approach to predict the orientation of CRISPR arrays. bioRxiv, 2025. doi: 10.1101/2025.05.09.653049.

41. Ling Deng, Roger A. Garrett, Shiraz A. Shah, Xu Peng, and Qunxin She. A novel interference mechanism by a type IIIB CRISPR-Cmr module in Sulfolobus. Molecular Microbiology, 87(5): 1088–1099, 2013. doi: 10.1111/mmi.12152.

42. Sonali Majumdar, Peng Zhao, Neil T. Pfister, Mark Compton, Sara Olson, Claiborne V.C. Glover, Lance Wells, Brenton R. Graveley, Rebecca M. Terns, and Michael P. Terns. Three CRISPR-Cas immune effector complexes coexist in Pyrococcus furiosus. RNA, 21(6): 1147–1158, 2015. doi: 10.1261/rna.049130.114.

43. Elena Vialetto, Yanying Yu, Scott P. Collins, Katharina G. Wandera, Lars Barquist, and Chase L. Beisel. A target expression threshold dictates invader defense and prevents autoimmunity by CRISPR-Cas13. Cell Host & Microbe, 30(8):1151–1162.e6, 2022. doi: 10.1016/j.chom.2022.05.013.

44. Shally R. Margolis and Alexander J. Meeske. Crosstalk between three CRISPR-Cas types enables primed type VI-A adaptation in Listeria seeligeri. Cell Host & Microbe, 33(9): 1550–1560.e4, 2025. doi: 10.1016/j.chom.2025.05.020.

