## Supplementary Material for "The complexity of multiple CRISPR arrays in strains with (co-occurring) CRISPR systems"

**Supplementary Table S1. CRISPR array per Cas locus statistics by Cas type, number of Cas loci with associated same repeat arrays and more.** We show the average, median and maximum number of arrays per Cas locus and the total locus count for the respective Cas types. In the second half of the table, 'same repeat' gives the amount of Cas loci which have at least one associated array that has the same consensus repeat as another array within the same genome. Then, each of the following columns gives the counts of Cas loci with associated arrays that are in a genome with a 'partner' array (not necessarily belonging to the same Cas locus), that *additionally* to having the same consensus repeat has some spacer content similarity. '+ Sp. Overlap' additionally requires some amount of spacer overlap (not necessarily the same spacer order). '+ = Sp.' additionally requires exactly the same spacer content (not necessarily the same spacer order). '+ Sp. Subset' additionally requires the spacer content to be contained in the other array (not necessarily the same spacer order). '+ = Last Sp.' additionally requires arrays to have exactly the same last spacer (according to the predicted orientation). All of the counts ignore spacer orientation, i.e. spacer and reverse complement are considered the same. Note that the 'partner' arrays are only part of the same genome and not required to belong to the same Cas locus or Cas type. Orphan arrays are excluded for this table. Supplementary Table S2 gives the same statistics in terms of array numbers instead of Cas loci.

| Cas Type | Average | Median | Max | Locus Count | Same Repeat | + Sp. Overlap | + = Sp. | + Sp. subset | + = Last Sp. |
|---|---|---|---|---|---|---|---|---|---|
| CAS-I-A | 1.55 | 1 | 5 | 149 | 71 | 4 | 0 | 0 | 0 |
| CAS-I-B | 1.46 | 1 | 9 | 2981 | 638 | 61 | 4 | 5 | 8 |
| CAS-I-C | 1.37 | 1 | 12 | 2824 | 286 | 37 | 8 | 13 | 13 |
| CAS-I-D | 1.74 | 2 | 8 | 363 | 124 | 6 | 1 | 1 | 1 |
| CAS-I-E | 1.73 | 2 | 12 | 10859 | 2907 | 59 | 10 | 11 | 13 |
| CAS-I-F | 1.70 | 2 | 8 | 3021 | 824 | 30 | 1 | 2 | 5 |
| CAS-I-U | 1.41 | 1 | 8 | 469 | 59 | 2 | 0 | 1 | 0 |
| CAS-II-A | 1.07 | 1 | 5 | 2884 | 182 | 17 | 3 | 3 | 3 |
| CAS-II-B | 1.21 | 1 | 11 | 147 | 18 | 4 | 4 | 4 | 4 |
| CAS-II-C | 1.20 | 1 | 10 | 3007 | 150 | 38 | 27 | 27 | 27 |
| CAS-III-A | 1.97 | 2 | 11 | 2507 | 1257 | 423 | 15 | 146 | 195 |
| CAS-III-B | 1.72 | 1 | 10 | 930 | 283 | 22 | 1 | 1 | 1 |
| CAS-III-C | 1.58 | 1 | 4 | 57 | 21 | 1 | 0 | 0 | 0 |
| CAS-III-D | 1.82 | 1 | 9 | 560 | 171 | 9 | 0 | 2 | 3 |
| CAS-III-E | 2.00 | 2 | 2 | 1 | 1 | 0 | 0 | 0 | 0 |
| CAS-IV-A | 1.17 | 1 | 3 | 123 | 4 | 1 | 0 | 0 | 0 |
| CAS-V-A | 1.26 | 1 | 23 | 4189 | 511 | 61 | 15 | 17 | 24 |
| CAS-V-B | 1.13 | 1 | 4 | 192 | 8 | 2 | 1 | 1 | 1 |
| CAS-V-F | 1.39 | 1 | 6 | 1436 | 210 | 27 | 2 | 8 | 10 |
| CAS-VI-A | 1.45 | 1 | 4 | 20 | 2 | 0 | 0 | 0 | 0 |
| CAS-VI-B | 1.31 | 1 | 11 | 7658 | 797 | 48 | 16 | 18 | 25 |
| CAS-VI-C | 1.22 | 1 | 9 | 383 | 99 | 3 | 0 | 0 | 0 |
| Overall | 1.48 | 1.0 | 23 | 44760 | 8623 | 855 | 108 | 260 | 333 |

**Supplementary Table S2. Locus and array count per Cas type, number of arrays with same repeat and more.** Besides the count of Cas loci and array count of all Cas types, we show the number of arrays which have the same consensus repeat as at least one other array in the same genome (as is reported in terms of Cas loci in Supplementary Table S1). Moreover, we show the number of arrays with same consensus repeat and additional spacer similarity (with at least one other array in the same genome) described in the caption of Supplementary Table S1.

| Cas Type | Locus Count | Array Count | Same Repeat | + Sp. Overlap | + = Sp. | + Sp. Subset | + = Last Sp. |
|---|---|---|---|---|---|---|---|
| CAS-I-A | 149 | 231 | 113 | 8 | 0 | 0 | 0 |
| CAS-I-B | 2981 | 4364 | 1156 | 120 | 13 | 16 | 26 |
| CAS-I-C | 2824 | 3856 | 598 | 72 | 16 | 23 | 27 |
| CAS-I-D | 363 | 630 | 219 | 12 | 2 | 2 | 2 |
| CAS-I-E | 10859 | 18732 | 5931 | 116 | 14 | 15 | 24 |
| CAS-I-F | 3021 | 5148 | 1500 | 47 | 1 | 2 | 8 |
| CAS-I-U | 469 | 661 | 121 | 4 | 0 | 1 | 0 |
| CAS-II-A | 2884 | 3075 | 243 | 26 | 3 | 3 | 3 |
| CAS-II-B | 147 | 178 | 32 | 4 | 4 | 4 | 4 |
| CAS-II-C | 3007 | 3621 | 283 | 71 | 51 | 51 | 52 |
| CAS-III-A | 2507 | 4932 | 3162 | 1229 | 32 | 179 | 431 |
| CAS-III-B | 930 | 1601 | 530 | 32 | 1 | 1 | 1 |
| CAS-III-C | 57 | 90 | 31 | 1 | 0 | 0 | 0 |
| CAS-III-D | 560 | 1020 | 324 | 14 | 0 | 3 | 6 |
| CAS-III-E | 1 | 2 | 2 | 0 | 0 | 0 | 0 |
| CAS-IV-A | 123 | 144 | 7 | 2 | 0 | 0 | 0 |
| CAS-V-A | 4189 | 5277 | 709 | 79 | 21 | 23 | 32 |
| CAS-V-B | 192 | 217 | 14 | 5 | 4 | 4 | 4 |
| CAS-V-F | 1436 | 1998 | 309 | 48 | 3 | 9 | 17 |
| CAS-VI-A | 20 | 29 | 3 | 0 | 0 | 0 | 0 |
| CAS-VI-B | 7658 | 10020 | 1158 | 74 | 27 | 29 | 38 |
| CAS-VI-C | 383 | 466 | 112 | 3 | 0 | 0 | 0 |
| Overall | 44760 | 66292 | 16557 | 1967 | 192 | 365 | 675 |

**Supplementary Table S3. CRISPR array length (the number of spacers in the array) statistics by Cas type.**

| Cas Type | Average | Median | Max | Array Count |
|---|---|---|---|---|
| CAS-I-A | 51.49 | 42 | 409 | 231 |
| CAS-I-B | 29.13 | 18 | 587 | 4364 |
| CAS-I-C | 25.26 | 15 | 318 | 3856 |
| CAS-I-D | 42.93 | 25 | 385 | 630 |
| CAS-I-E | 16.78 | 11 | 246 | 18732 |
| CAS-I-F | 21.58 | 14.5 | 317 | 5148 |
| CAS-I-U | 25.06 | 15 | 587 | 661 |
| CAS-II-A | 15.93 | 12 | 158 | 3075 |
| CAS-II-B | 17.76 | 11 | 71 | 178 |
| CAS-II-C | 11.92 | 5 | 184 | 3621 |
| CAS-III-A | 15.71 | 14 | 276 | 4932 |
| CAS-III-B | 19.83 | 11 | 359 | 1601 |
| CAS-III-C | 27.39 | 17 | 131 | 90 |
| CAS-III-D | 19.89 | 9 | 233 | 1020 |
| CAS-III-E | 6.50 | 6.5 | 7 | 2 |
| CAS-IV-A | 10.51 | 10 | 69 | 144 |
| CAS-V-A | 8.24 | 4 | 204 | 5277 |
| CAS-V-B | 5.68 | 3 | 48 | 217 |
| CAS-V-F | 8.24 | 5 | 102 | 1998 |
| CAS-VI-A | 10.14 | 4 | 69 | 29 |
| CAS-VI-B | 6.83 | 4 | 300 | 10020 |
| CAS-VI-C | 5.35 | 3 | 66 | 466 |
| Orphan | 11.25 | 7 | 242 | 16689 |
| Overall | 15.11 | 9 | 587 | 82981 |

**Supplementary Table S4. CRISPR array length (the number of spacers in the array) statistics by Cas type broken down by 'Cas is close' label.** The 'Cas is close' is *yes* if the distance to its respective Cas locus is less than 10,000 bp and *no* otherwise. Orphan arrays are excluded for this table.

| Cas Type | Cas is close | Average | Median | Max | Array Count |
|---|---|---|---|---|---|
| CAS-I-A | yes | 52.11 | 39 | 192 | 169 |
| | no | 49.79 | 44.5 | 409 | 62 |
| CAS-I-B | yes | 32.03 | 22 | 587 | 3483 |
| | no | 17.68 | 7 | 374 | 881 |
| CAS-I-C | yes | 27.39 | 17 | 318 | 3293 |
| | no | 12.82 | 6 | 181 | 563 |
| CAS-I-D | yes | 48.71 | 33 | 164 | 484 |
| | no | 23.77 | 8 | 385 | 146 |
| CAS-I-E | yes | 18.10 | 12 | 246 | 14949 |
| | no | 11.54 | 8 | 184 | 3783 |
| CAS-I-F | yes | 22.50 | 16 | 317 | 4765 |
| | no | 10.07 | 6 | 97 | 383 |
| CAS-I-U | yes | 27.69 | 19 | 587 | 549 |
| | no | 12.14 | 4 | 146 | 112 |
| CAS-II-A | yes | 16.28 | 12 | 158 | 2969 |
| | no | 6.25 | 3 | 41 | 106 |
| CAS-II-B | yes | 20.47 | 13 | 71 | 131 |
| | no | 10.21 | 5 | 50 | 47 |
| CAS-II-C | yes | 17.41 | 12 | 184 | 2003 |
| | no | 5.12 | 4 | 145 | 1618 |
| CAS-III-A | yes | 15.98 | 15 | 276 | 4529 |
| | no | 12.66 | 7.0 | 165 | 403 |
| CAS-III-B | yes | 19.76 | 12 | 179 | 1265 |
| | no | 20.11 | 8 | 359 | 336 |
| CAS-III-C | yes | 29.51 | 17 | 131 | 63 |
| | no | 17.01 | 8 | 233 | 319 |
| CAS-III-D | yes | 21.20 | 11 | 198 | 701 |
| | no | 22.44 | 8 | 86 | 27 |
| CAS-III-E | yes | 6.5 | 6.5 | 7 | 2 |
| | no | - | - | - | 0 |
| CAS-IV-A | yes | 11.24 | 10 | 69 | 116 |
| | no | 7.5 | 4 | 25 | 28 |
| CAS-V-A | yes | 15.14 | 10 | 173 | 359 |
| | no | 7.74 | 4 | 204 | 4918 |
| CAS-V-B | yes | 14.35 | 7 | 48 | 17 |
| | no | 4.94 | 3 | 36 | 200 |
| CAS-V-F | yes | 11.93 | 7 | 102 | 103 |
| | no | 8.03 | 5 | 94 | 1895 |
| CAS-VI-A | yes | 8.00 | 4 | 23 | 5 |
| | no | 10.58 | 4 | 69 | 24 |
| CAS-VI-B | yes | 11.88 | 7 | 213 | 418 |
| | no | 6.61 | 4 | 300 | 9602 |
| CAS-VI-C | yes | 3.90 | 2.5 | 11 | 10 |
| | no | 5.38 | 3 | 66 | 456 |
| Overall | yes | 20.81 | 14 | 587 | 40383 |
| | no | 8.73 | 4 | 409 | 25909 |

**Supplementary Table S5. Underlying table for the diagonal of Figure 1 in the main manuscript.** The table shows the fold-change aloneness, expected and observed frequencies and counts for each Cas type as well as the p-value of the bionomial test. The last column shows significance, i.e. *p < 0.05, **p < 0.01, ***p < 0.001, ****p < 0.0001, if there are more than 50 observed or expected observations of a type.

| Cas Type | Fold-change | Ratio | Obs. Freq | Exp. Freq | Obs. Count | Exp. Count | p-value | Signif. |
|---|---|---|---|---|---|---|---|---|
| CAS-I-A | -1.3374 | 0.7477 | 0.00083 | 0.00112 | 15 | 20.06 | 0.1533 | |
| CAS-I-B | 1.7150 | 1.7150 | 0.02914 | 0.01699 | 524 | 305.54 | 5.94E-14 | **** |
| CAS-I-C | 1.4737 | 1.4737 | 0.02308 | 0.01566 | 415 | 281.61 | <1.00E-15 | **** |
| CAS-I-D | 1.3264 | 1.3264 | 0.00378 | 0.00285 | 68 | 51.27 | 0.0103 | |
| CAS-I-E | 2.5939 | 2.5939 | 0.33343 | 0.12854 | 5996 | 2311.57 | <1.00E-15 | **** |
| CAS-I-F | 3.3511 | 3.3511 | 0.07446 | 0.02222 | 1339 | 399.57 | 1.12E-12 | **** |
| CAS-I-U | 1.3027 | 1.3027 | 0.00328 | 0.00252 | 59 | 45.29 | 0.0207 | * |
| CAS-II-A | -1.1843 | 0.8444 | 0.00945 | 0.01120 | 170 | 201.33 | 0.0128 | * |
| CAS-II-C | -1.7100 | 0.5848 | 0.01001 | 0.01712 | 180 | 307.81 | <1.00E-15 | **** |
| CAS-III-A | -1.5883 | 0.6296 | 0.01462 | 0.02323 | 263 | 417.73 | <1.00E-15 | **** |
| CAS-III-B | 1.2164 | 1.2164 | 0.00823 | 0.00677 | 148 | 121.67 | 0.0088 | ** |
| CAS-III-D | 1.1140 | 1.1140 | 0.00462 | 0.00414 | 83 | 74.51 | 0.1484 | |
| CAS-V-A | -1.2758 | 0.7838 | 0.01946 | 0.02483 | 350 | 446.56 | 9.09E-07 | **** |
| CAS-V-B | -2.3441 | 0.4266 | 0.00050 | 0.00117 | 9 | 21.10 | 0.0026 | |
| CAS-V-F | -1.7241 | 0.5800 | 0.00473 | 0.00815 | 85 | 146.54 | 2.17E-08 | **** |
| CAS-VI-B | -1.1444 | 0.8738 | 0.05210 | 0.05963 | 937 | 1072.37 | 7.43E-06 | **** |
| CAS-VI-C | -2.2904 | 0.4366 | 0.00117 | 0.00267 | 21 | 48.10 | 9.05E-06 | **** |

**Supplementary Table S6. Underlying table for all off-diagonal entries in Figure 1 in the main manuscript.** The table shows the fold-change co-occurrence, expected and observed frequencies and counts for each Cas type as well as the p-value of the bionomial test. The last column shows significance, i.e. *p < 0.05, **p < 0.01, ***p < 0.001, ****p < 0.0001, if there are more than 50 observed or expected observations of a type pair. We omit pairs without any observed occurrences.

| 1. Cas Type | 2. Cas Type | Fold-change | Ratio | Obs. Freq. | Exp. Freq. | Obs. Count | Exp. Count | p-value | Signif. |
|---|---|---|---|---|---|---|---|---|---|
| CAS-I-A | CAS-I-B | -2.0008 | 0.4998 | 5.56E-05 | 1.11E-04 | 1 | 2.00 | 0.4057 | |
| CAS-I-A | CAS-I-C | -1.8443 | 0.5422 | 5.56E-05 | 1.03E-04 | 1 | 1.84 | 0.4498 | |
| CAS-I-A | CAS-I-D | 2.9786 | 2.9786 | 5.56E-05 | 1.87E-05 | 1 | 0.34 | 0.0452 | |
| CAS-I-A | CAS-I-E | -7.5700 | 0.1321 | 1.11E-04 | 8.42E-04 | 2 | 15.14 | 3.46E-05 | |
| CAS-I-A | CAS-III-A | -1.368 | 0.7311 | 1.11E-04 | 1.52E-04 | 2 | 2.74 | 0.4849 | |
| CAS-I-A | CAS-III-B | 11.2950 | 11.2950 | 5.00E-04 | 4.43E-05 | 9 | 0.80 | 1.38E-08 | |
| CAS-I-A | CAS-III-D | 22.5448 | 22.5448 | 6.12E-04 | 2.71E-05 | 11 | 0.49 | 7.40E-13 | |
| CAS-I-A | CAS-V-A | 1.0258 | 1.0258 | 1.67E-04 | 1.63E-04 | 3 | 2.92 | 0.3358 | |
| CAS-I-A | CAS-V-F | 2.0841 | 2.0841 | 1.11E-04 | 5.34E-05 | 2 | 0.96 | 0.0730 | |
| CAS-I-A | CAS-VI-B | -1.0032 | 0.9968 | 3.89E-04 | 3.91E-04 | 7 | 7.023 | 0.5953 | |
| CAS-I-B | CAS-I-C | 1.1749 | 1.1749 | 1.84E-03 | 1.56E-03 | 33 | 28.09 | 0.1534 | |
| CAS-I-B | CAS-I-D | 1.3690 | 1.3690 | 3.89E-04 | 2.84E-04 | 7 | 5.11 | 0.1454 | |
| CAS-I-B | CAS-I-E | -6.4061 | 0.1561 | 2.00E-03 | 1.28E-02 | 36 | 230.56 | <1.00E-15 | **** |
| CAS-I-B | CAS-I-F | -9.9602 | 0.1004 | 2.22E-04 | 2.22E-03 | 4 | 39.85 | 5.53E-13 | |
| CAS-I-B | CAS-I-U | 2.4351 | 2.4351 | 6.12E-04 | 2.51E-04 | 11 | 4.52 | 0.0025 | |
| CAS-I-B | CAS-II-A | 1.3944 | 1.3944 | 1.56E-03 | 1.12E-03 | 28 | 20.08 | 0.0357 | |
| CAS-I-B | CAS-II-C | 1.1726 | 1.1726 | 2.00E-03 | 1.71E-03 | 36 | 30.70 | 0.1478 | |
| CAS-I-B | CAS-III-A | 1.2000 | 1.2000 | 2.78E-03 | 2.32E-03 | 50 | 41.67 | 0.0884 | |
| CAS-I-B | CAS-III-B | 5.1911 | 5.1911 | 3.50E-03 | 6.75E-04 | 63 | 12.14 | 8.11E-13 | **** |
| CAS-I-B | CAS-III-D | 3.6332 | 3.6332 | 1.50E-03 | 4.13E-04 | 27 | 7.43 | 6.32E-09 | |
| CAS-I-B | CAS-V-A | 1.8634 | 1.8634 | 4.62E-03 | 2.48E-03 | 83 | 44.54 | 8.50E-08 | **** |
| CAS-I-B | CAS-V-B | -2.1044 | 0.4752 | 5.56E-05 | 1.17E-04 | 1 | 2.10 | 0.3785 | |
| CAS-I-B | CAS-V-F | -1.0441 | 0.9578 | 7.79E-04 | 8.13E-04 | 14 | 14.62 | 0.5054 | |
| CAS-I-B | CAS-VI-B | 1.1967 | 1.1967 | 7.12E-03 | 5.95E-03 | 128 | 106.96 | 0.0207 | * |
| CAS-I-B | CAS-VI-C | -1.5992 | 0.6253 | 1.67E-04 | 2.67E-04 | 3 | 4.80 | 0.2946 | |
| CAS-I-C | CAS-I-D | -2.3563 | 0.4244 | 1.11E-04 | 2.62E-04 | 2 | 4.71 | 0.1510 | |
| CAS-I-C | CAS-I-E | -4.1667 | 0.2400 | 2.84E-03 | 1.18E-02 | 51 | 212.50 | <1.00E-15 | **** |
| CAS-I-C | CAS-I-F | 1.6062 | 1.6062 | 3.28E-03 | 2.04E-03 | 59 | 36.73 | 0.0003 | *** |
| CAS-I-C | CAS-I-U | 3.6028 | 3.6028 | 8.34E-04 | 2.32E-04 | 15 | 4.16 | 7.95E-06 | |
| CAS-I-C | CAS-II-A | 17.8843 | 17.8843 | 1.84E-02 | 1.03E-03 | 331 | 18.51 | <1.00E-15 | **** |
| CAS-I-C | CAS-II-C | 2.3678 | 2.3678 | 3.73E-03 | 1.57E-03 | 67 | 28.30 | 1.74E-10 | **** |
| CAS-I-C | CAS-III-A | -1.5361 | 0.6510 | 1.39E-03 | 2.14E-03 | 25 | 38.40 | 0.0142 | |
| CAS-I-C | CAS-III-B | 1.7880 | 1.7880 | 1.11E-03 | 6.22E-04 | 20 | 11.19 | 0.0056 | |
| CAS-I-C | CAS-III-D | 1.6060 | 1.6060 | 6.12E-04 | 3.81E-04 | 11 | 6.85 | 0.4468 | |
| CAS-I-C | CAS-V-A | 2.4116 | 2.4116 | 5.51E-03 | 2.28E-03 | 99 | 41.05 | 1.00E-13 | **** |
| CAS-I-C | CAS-V-F | -2.6940 | 0.3712 | 2.78E-04 | 7.49E-04 | 5 | 13.47 | 0.0079 | |
| CAS-I-C | CAS-VI-A | 7.9752 | 7.9752 | 1.11E-04 | 1.39E-05 | 2 | 0.25 | 0.0022 | |
| CAS-I-C | CAS-VI-B | 1.0448 | 1.0448 | 5.73E-03 | 5.48E-03 | 103 | 98.58 | 0.3054 | |
| CAS-I-C | CAS-VI-C | -4.4209 | 0.2262 | 5.56E-05 | 2.46E-04 | 1 | 4.42 | 0.0651 | |
| CAS-I-D | CAS-I-E | -38.7597 | 0.0258 | 5.56E-05 | 2.15E-03 | 1 | 38.69 | <1.00E-15 | |
| CAS-I-D | CAS-I-F | -6.6890 | 0.1495 | 5.56E-05 | 3.72E-04 | 1 | 6.69 | 0.0096 | |
| CAS-I-D | CAS-II-C | -1.2878 | 0.7765 | 2.22E-04 | 2.86E-04 | 4 | 5.15 | 0.4143 | |
| CAS-I-D | CAS-III-B | 3.9288 | 3.9288 | 4.45E-04 | 1.13E-04 | 8 | 2.04 | 0.0003 | |
| CAS-I-D | CAS-III-D | 1.6040 | 1.6040 | 1.11E-04 | 6.93E-05 | 2 | 1.25 | 0.1308 | |
| CAS-I-D | CAS-V-A | 3.7466 | 3.7466 | 1.56E-03 | 4.16E-04 | 28 | 7.47 | 1.81E-09 | |
| CAS-I-D | CAS-V-F | -2.4522 | 0.4078 | 5.56E-05 | 1.36E-04 | 1 | 2.45 | 0.2972 | |
| CAS-I-D | CAS-VI-B | -1.7947 | 0.5572 | 5.56E-04 | 9.98E-04 | 10 | 17.95 | 0.0311 | |
| CAS-I-D | CAS-VI-C | 1.2423 | 1.2423 | 5.56E-05 | 4.48E-05 | 1 | 0.80 | 0.1930 | |
| CAS-I-E | CAS-I-F | -2.4120 | 0.4146 | 6.95E-03 | 1.68E-02 | 125 | 301.52 | <1.00E-15 | **** |
| CAS-I-E | CAS-I-U | -1.1786 | 0.8485 | 1.61E-03 | 1.90E-03 | 29 | 34.18 | 0.2146 | |
| CAS-I-E | CAS-II-A | -2.0255 | 0.4937 | 4.17E-03 | 8.45E-03 | 75 | 151.92 | 2.92E-12 | **** |
| CAS-I-E | CAS-II-C | -4.8379 | 0.2067 | 2.67E-03 | 1.29E-02 | 48 | 232.27 | <1.00E-15 | **** |
| CAS-I-E | CAS-III-A | -15.7729 | 0.0634 | 1.11E-03 | 1.75E-02 | 20 | 315.22 | <1.00E-15 | **** |
| CAS-I-E | CAS-III-B | -3.2787 | 0.3050 | 1.56E-03 | 5.11E-03 | 28 | 91.82 | 1.00E-15 | **** |
| CAS-I-E | CAS-III-D | -2.8114 | 0.3557 | 1.11E-03 | 3.13E-03 | 20 | 56.22 | 2.31E-08 | **** |
| CAS-I-E | CAS-V-A | -1.2812 | 0.7805 | 1.46E-02 | 1.87E-02 | 263 | 336.98 | 1.39E-05 | **** |
| CAS-I-E | CAS-V-B | 3.4549 | 3.4549 | 3.06E-03 | 8.85E-04 | 55 | 15.92 | 1.17E-12 | **** |

| 1. Cas Type | 2. Cas Type | Fold-change | Ratio | Obs. Freq. | Exp. Freq. | Obs. Count | Exp. Count | p-value | Signif. |
|---|---|---|---|---|---|---|---|---|---|
| CAS-I-E | CAS-V-F | -4.2535 | 0.2351 | 1.45E-03 | 6.15E-03 | 26 | 110.58 | <1.00E-15 | **** |
| CAS-I-E | CAS-VI-A | -2.0585 | 0.4858 | 5.56E-05 | 1.14E-04 | 1 | 2.06 | 0.3904 | |
| CAS-I-E | CAS-VI-B | -1.4049 | 0.7118 | 3.20E-02 | 4.50E-02 | 576 | 809.22 | <1.00E-15 | **** |
| CAS-I-E | CAS-VI-C | -1.3961 | 0.7163 | 1.45E-03 | 2.02E-03 | 26 | 36.30 | 0.0464 | |
| CAS-I-F | CAS-I-U | -2.9533 | 0.3386 | 1.11E-04 | 3.29E-04 | 2 | 5.91 | 0.0662 | |
| CAS-I-F | CAS-II-C | -2.1133 | 0.4732 | 1.06E-03 | 2.23E-03 | 19 | 40.15 | 0.0002 | |
| CAS-I-F | CAS-III-A | -9.0827 | 0.1101 | 3.34E-04 | 3.03E-03 | 6 | 54.49 | 1.00E-15 | **** |
| CAS-I-F | CAS-III-B | -2.6448 | 0.3781 | 3.34E-04 | 8.83E-04 | 6 | 15.87 | 0.0043 | |
| CAS-I-F | CAS-III-D | -2.4295 | 0.4116 | 2.22E-04 | 5.40E-04 | 4 | 9.72 | 0.035 | |
| CAS-I-F | CAS-V-A | 2.0773 | 2.0773 | 6.73E-03 | 3.24E-03 | 121 | 58.25 | <1.00E-15 | **** |
| CAS-I-F | CAS-V-B | 1.8170 | 1.8170 | 2.78E-04 | 1.53E-04 | 5 | 2.75 | 0.0610 | |
| CAS-I-F | CAS-V-F | -2.1240 | 0.4708 | 5.00E-04 | 1.06E-03 | 9 | 19.11 | 0.0083 | |
| CAS-I-F | CAS-VI-B | 1.3798 | 1.3798 | 1.07E-02 | 7.78E-03 | 193 | 139.88 | 8.00E-06 | **** |
| CAS-I-F | CAS-VI-C | 4.6223 | 4.6223 | 1.61E-03 | 3.49E-04 | 29 | 6.27 | 7.12E-12 | |
| CAS-I-U | CAS-II-A | -1.4883 | 0.6719 | 1.11E-04 | 1.66E-04 | 2 | 2.98 | 0.4284 | |
| CAS-I-U | CAS-II-C | 1.7579 | 1.7579 | 4.45E-04 | 2.53E-04 | 8 | 4.55 | 0.0426 | |
| CAS-I-U | CAS-III-A | -2.0585 | 0.4858 | 1.67E-04 | 3.43E-04 | 3 | 6.18 | 0.1361 | |
| CAS-I-U | CAS-III-B | 3.3354 | 3.3354 | 3.36E-04 | 1.00E-04 | 6 | 1.80 | 0.0026 | |
| CAS-I-U | CAS-III-D | 3.6313 | 3.6313 | 2.22E-04 | 6.13E-05 | 4 | 1.10 | 0.0055 | |
| CAS-I-U | CAS-V-A | 1.2117 | 1.2117 | 4.45E-04 | 3.67E-04 | 8 | 6.60 | 0.2207 | |
| CAS-I-U | CAS-V-F | -1.0833 | 0.9231 | 1.11E-04 | 1.20E-04 | 2 | 2.17 | 0.6317 | |
| CAS-I-U | CAS-VI-B | 1.1353 | 1.1353 | 1.00E-03 | 8.82E-04 | 18 | 15.85 | 0.2456 | |
| CAS-II-A | CAS-II-C | -20.2429 | 0.0494 | 5.56E-05 | 1.12E-03 | 1 | 20.23 | 3.44E-08 | |
| CAS-II-A | CAS-III-A | 1.8577 | 1.8577 | 2.84E-03 | 1.53E-03 | 51 | 27.45 | 1.90E-05 | **** |
| CAS-II-A | CAS-III-B | -3.9984 | 0.2501 | 1.11E-04 | 4.45E-04 | 2 | 8.00 | 0.0138 | |
| CAS-II-A | CAS-III-D | -1.2241 | 0.8169 | 2.22E-04 | 2.72E-04 | 4 | 4.90 | 0.4588 | |
| CAS-II-A | CAS-V-A | 4.8383 | 4.8383 | 7.90E-03 | 1.63E-03 | 142 | 29.35 | <1.00E-15 | **** |
| CAS-II-A | CAS-V-F | 1.3498 | 1.3498 | 7.23E-04 | 5.36E-04 | 13 | 9.63 | 0.1101 | |
| CAS-II-A | CAS-VI-B | 2.6533 | 2.6533 | 1.04E-02 | 3.92E-03 | 187 | 70.48 | 1.34E-12 | **** |
| CAS-II-A | CAS-VI-C | -1.5805 | 0.6327 | 1.11E-04 | 1.76E-04 | 2 | 3.16 | 0.3880 | |
| CAS-II-C | CAS-III-A | 5.5748 | 5.5748 | 1.30E-02 | 2.33E-03 | 234 | 41.97 | 8.26E-13 | **** |
| CAS-II-C | CAS-III-B | -1.2226 | 0.8179 | 5.56E-04 | 6.80E-04 | 10 | 12.23 | 0.3239 | |
| CAS-II-C | CAS-III-D | 2.29386 | 2.9386 | 1.22E-03 | 4.16E-04 | 22 | 7.49 | 3.99E-06 | |
| CAS-II-C | CAS-V-A | 1.0474 | 1.0474 | 2.61E-03 | 2.50E-03 | 47 | 44.87 | 0.3395 | |
| CAS-II-C | CAS-V-F | 1.3582 | 1.3582 | 1.11E-03 | 8.19E-04 | 20 | 14.72 | 0.0719 | |
| CAS-II-C | CAS-VI-A | 3.6482 | 3.6482 | 5.56E-05 | 1.52E-05 | 1 | 0.27 | 0.0314 | |
| CAS-II-C | CAS-VI-B | 1.6797 | 1.6797 | 1.00E-02 | 5.99E-03 | 181 | 107.76 | 3.92E-11 | **** |
| CAS-II-C | CAS-VI-C | -4.8333 | 0.2069 | 5.56E-05 | 2.69E-04 | 1 | 4.83 | 0.0464 | |
| CAS-III-A | CAS-III-B | -41.1477 | 0.2411 | 2.22E-04 | 9.23E-04 | 4 | 16.59 | 0.0003 | |
| CAS-III-A | CAS-III-D | -2.5400 | 0.3937 | 2.22E-04 | 5.65E-04 | 4 | 10.16 | 0.0263 | |
| CAS-III-A | CAS-V-A | -3.0451 | 0.3284 | 1.11E-03 | 3.39E-03 | 20 | 60.90 | 1.02E-09 | **** |
| CAS-III-A | CAS-V-F | 5.1543 | 5.1543 | 5.73E-03 | 1.11E-03 | 103 | 19.98 | <1.00E-15 | **** |
| CAS-III-A | CAS-VI-B | 3.2208 | 3.2208 | 2.62E-02 | 8.13E-03 | 471 | 146.24 | 6.54E-13 | **** |
| CAS-III-A | CAS-VI-C | -6.5574 | 0.1525 | 5.56E-05 | 3.65E-04 | 1 | 6.56 | 0.0107 | |
| CAS-III-B | CAS-III-D | 1.6896 | 1.6896 | 2.78E-04 | 1.65E-04 | 5 | 2.96 | 0.0799 | |
| CAS-III-B | CAS-V-A | 1.2967 | 1.2967 | 1.2E-03 | 9.86E-04 | 23 | 17.74 | 0.0899 | |
| CAS-III-B | CAS-V-F | -1.1641 | 0.8590 | 2.78E-04 | 3.24E-04 | 5 | 5.82 | 0.4749 | |
| CAS-III-B | CAS-VI-B | -4.7326 | 0.2113 | 5.00E-04 | 2.37E-03 | 9 | 42.59 | 4.92E-10 | |
| CAS-III-B | CAS-VI-C | -1.9106 | 0.5234 | 5.56E-05 | 1.06E-04 | 1 | 1.91 | 0.4308 | |
| CAS-III-D | CAS-V-A | 1.1969 | 1.1969 | 7.23E-04 | 6.04E-04 | 13 | 10.86 | 0.2060 | |
| CAS-III-D | CAS-V-F | 1.1223 | 1.1223 | 2.22E-04 | 1.98E-04 | 4 | 3.56 | 0.2867 | |
| CAS-III-D | CAS-VI-B | -2.3714 | 0.4217 | 6.12E-04 | 1.45E-03 | 11 | 26.08 | 0.0007 | |
| CAS-V-A | CAS-V-B | 4.8774 | 4.8774 | 8.34E-04 | 1.71E-04 | 15 | 3.08 | 1.71E-07 | |
| CAS-V-A | CAS-V-F | 1.7788 | 1.7788 | 2.11E-03 | 1.19E-03 | 38 | 21.36 | 0.0004 | |
| CAS-V-A | CAS-VI-B | 1.8615 | 1.8615 | 1.62E-02 | 8.69E-03 | 291 | 156.33 | 9.17E-14 | **** |
| CAS-V-A | CAS-VI-C | 5.1342 | 5.1342 | 2.00E-03 | 3.90E-04 | 36 | 7.01 | 4.72E-13 | |
| CAS-V-B | CAS-V-F | 1.9817 | 1.9817 | 1.11E-04 | 5.61E-05 | 2 | 1.01 | 0.0820 | |
| CAS-V-B | CAS-VI-B | 1.3540 | 1.3540 | 5.56E-04 | 4.11E-04 | 10 | 7.39 | 0.1281 | |
| CAS-V-F | CAS-VI-A | 7.6630 | 7.6630 | 5.56E-05 | 7.26E-06 | 1 | 0.13 | 0.0078 | |
| CAS-V-F | CAS-VI-B | -1.0059 | 0.9941 | 2.84E-03 | 2.85E-03 | 51 | 51.30 | 0.5204 | |
| CAS-V-F | CAS-VI-C | -1.1505 | 0.8692 | 1.11E-04 | 1.28E-04 | 2 | 2.30 | 0.5958 | |
| CAS-VI-A | CAS-VI-B | 1.0472 | 1.0472 | 5.56E-05 | 5.31E-05 | 1 | 0.95 | 0.2477 | |

| 1. Cas Type | 2. Cas Type | Fold-change | Ratio | Obs. Freq. | Exp. Freq. | Obs. Count | Exp. Count | p-value | Signif. |
|---|---|---|---|---|---|---|---|---|---|
| CAS-VI-B | CAS-VI-C | 1.5441 | 1.5441 | 1.45E-03 | 9.36E-04 | 26 | 16.84 | 0.0136 | |

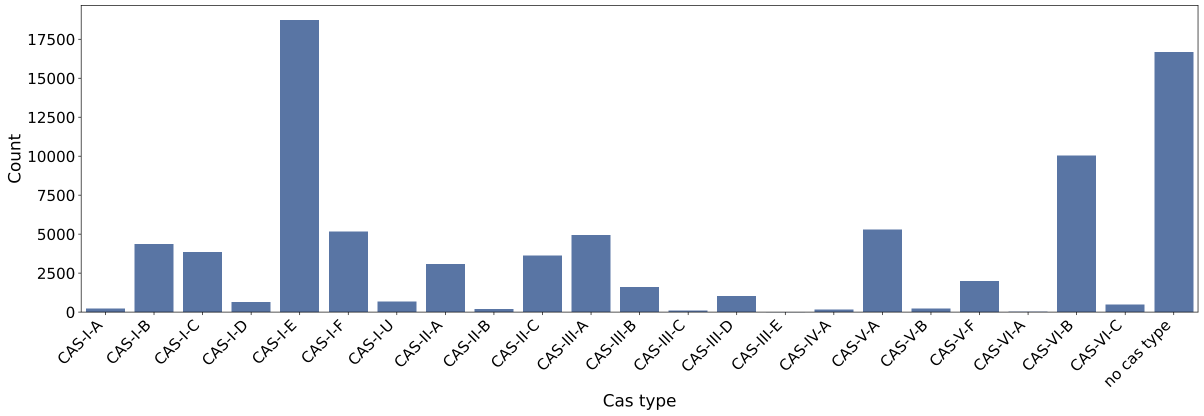


**Supplementary Fig. S1. Distribution of Cas types across arrays.** We show the distribution of Cas types of the arrays for the complete NCBI dataset.

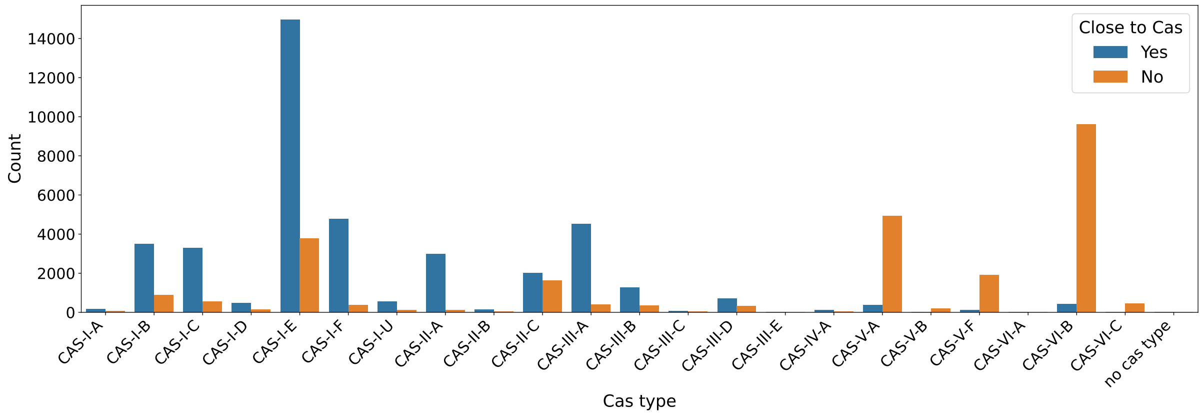


**Supplementary Fig. S2. Distribution of Cas types across arrays with 'Cas is close' label.** We break down the counts according to closeness of their Cas loci. We label a array to have a close Cas locus, if the Cas locus is less than 10000 bp removed from its Cas. Clearly, types V and VI tend to be far removed from their Cas loci and II-C shows a remarkable equal distribution between closeness and not.

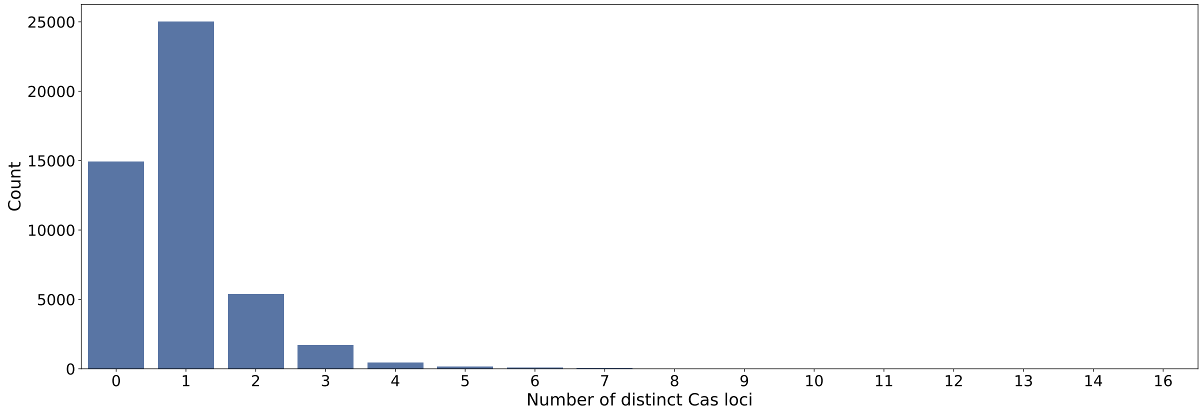


**Supplementary Fig. S3. Distribution of number of distinct Cas loci within genomes.** Here we show the distribution of the number of *distinct* Cas loci (not types!) per sample. 0 indicates that the array(s) within the genome are "orphan" arrays an do not contain *any* Cas locus. Since CRISPR arrays can only be orphans if no Cas locus exisits at all in the genome, there are no orphans in any other category than 0.

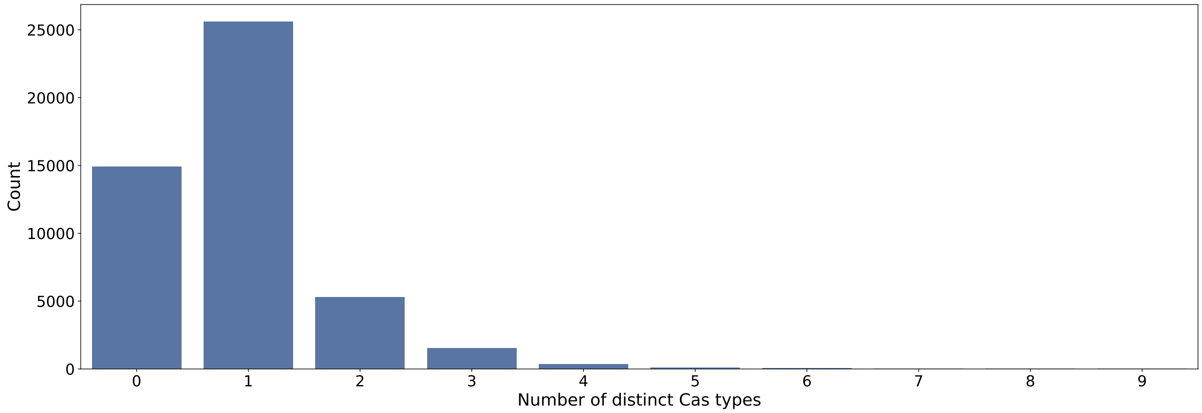


**Supplementary Fig. S4. Distribution of number of distinct Cas types within genomes.** Here a number of distinct Cas types of 0 means that the arrays within the genome are "orphan" arrays and do not contain *any* Cas loci. Since CRISPR arrays can only be orphans if no Cas locus exisits at all in the genome, there are no orphans in any other category than 0.

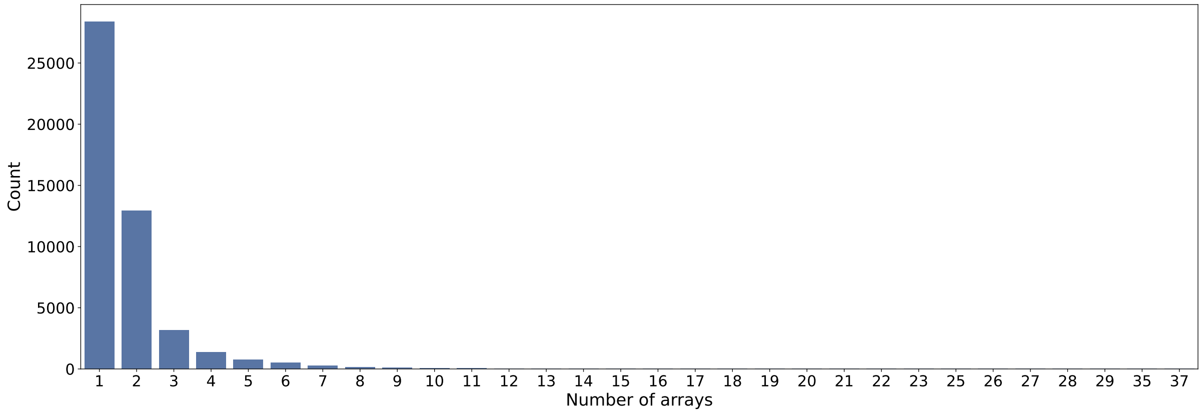


**Supplementary Fig. S5. Distribution of number of arrays within genomes.** We show all numbers on the x-axis for which at least one genome exists.

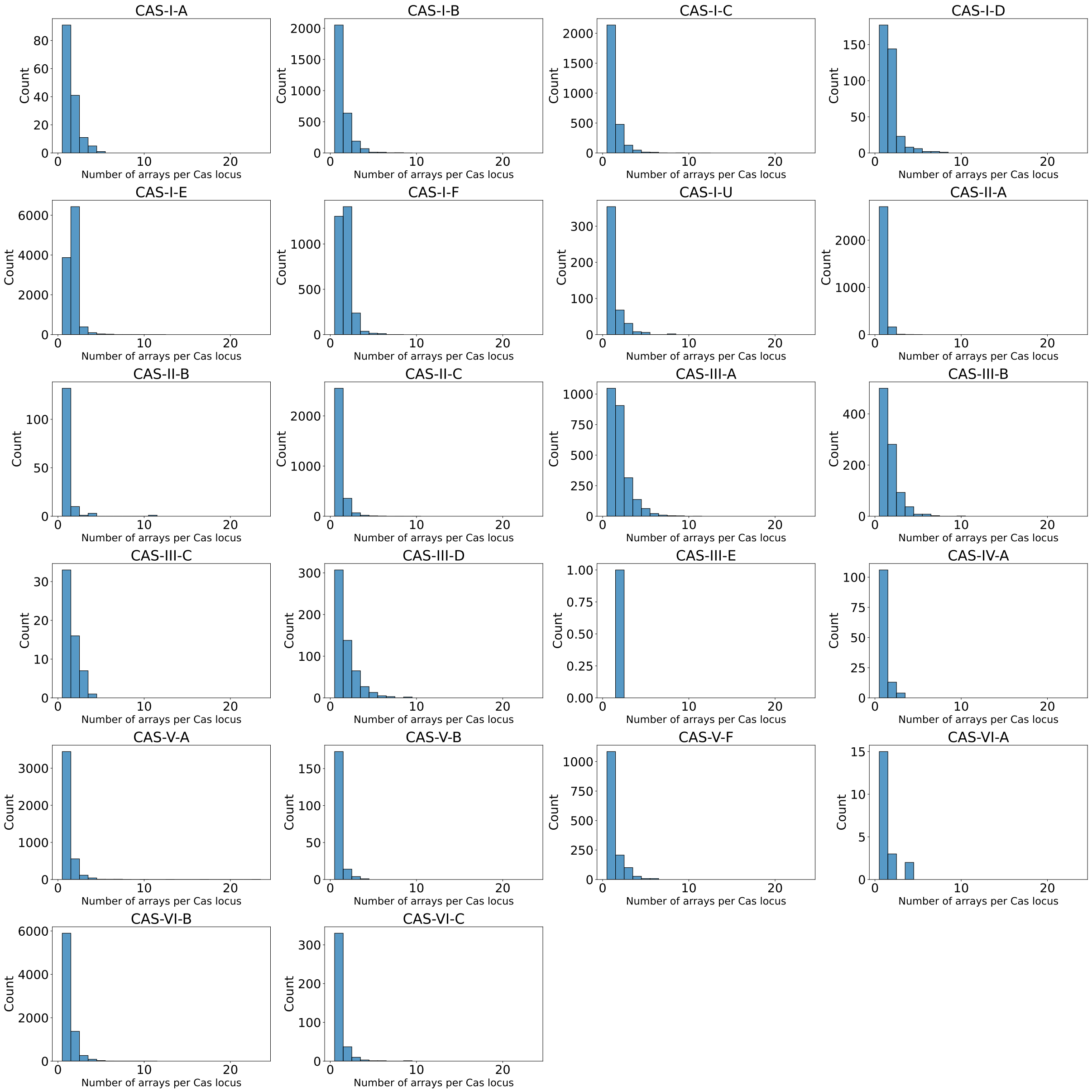


**Supplementary Fig. S6. Distribution of number of arrays per Cas locus for different Cas types.** We show distribution of the number of arrays belonging to a particular Cas locus. Type I-E, I-F and III-A tend to favor two arrays per Cas locus. Otherwise the distributions vary but are largely consistent.

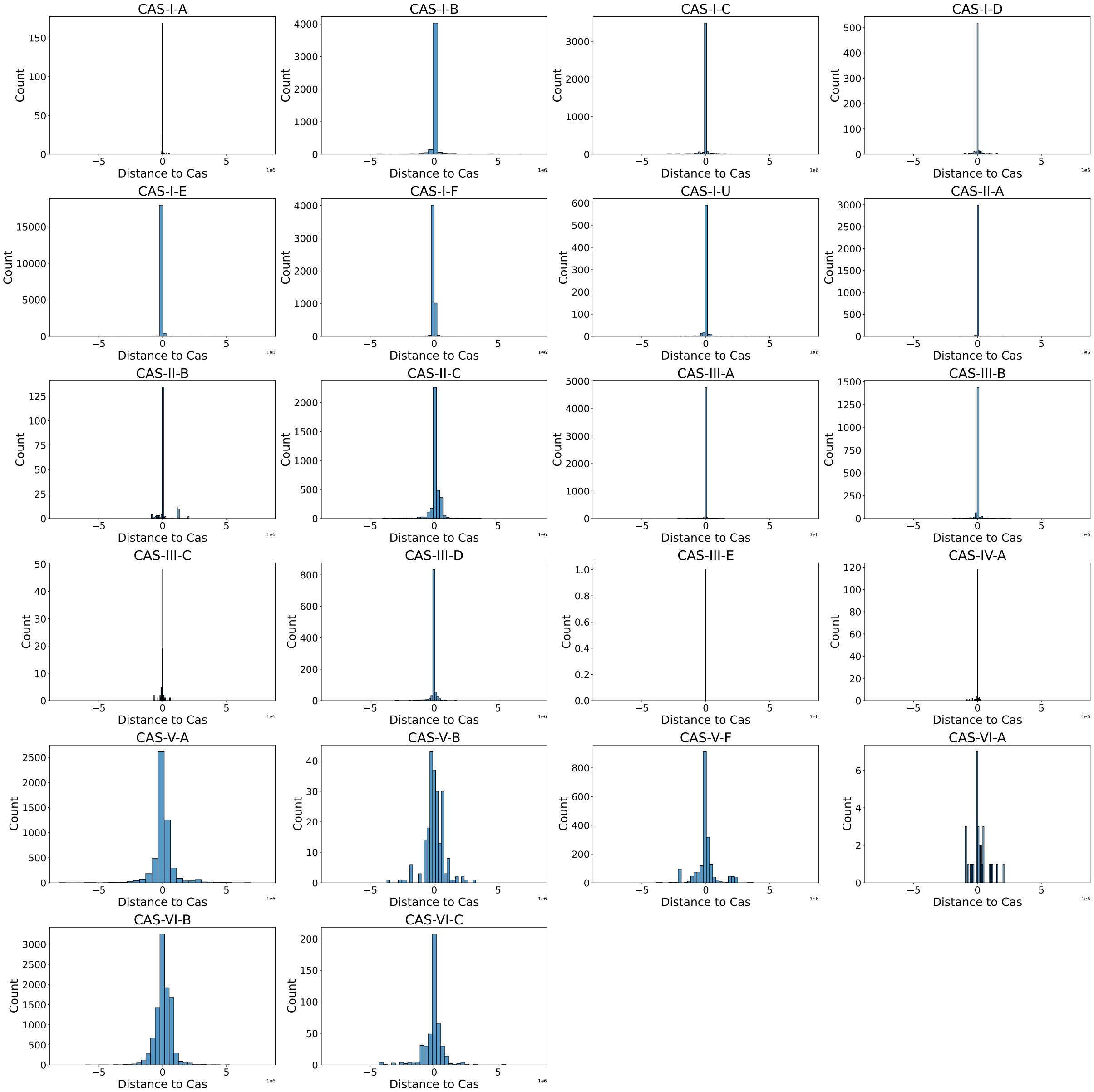


**Supplementary Fig. S7. Directional distances of CRISPR arrays to their respective Cas locus.** This figure clearly shows that types I, II and III are very concentrated around their Cas locus. Types V and VI however tend to be much further from their Cas loci. We excluded types with very few samples. Note that the x-axis is scaled in units of $10^6$ base pairs (bp).

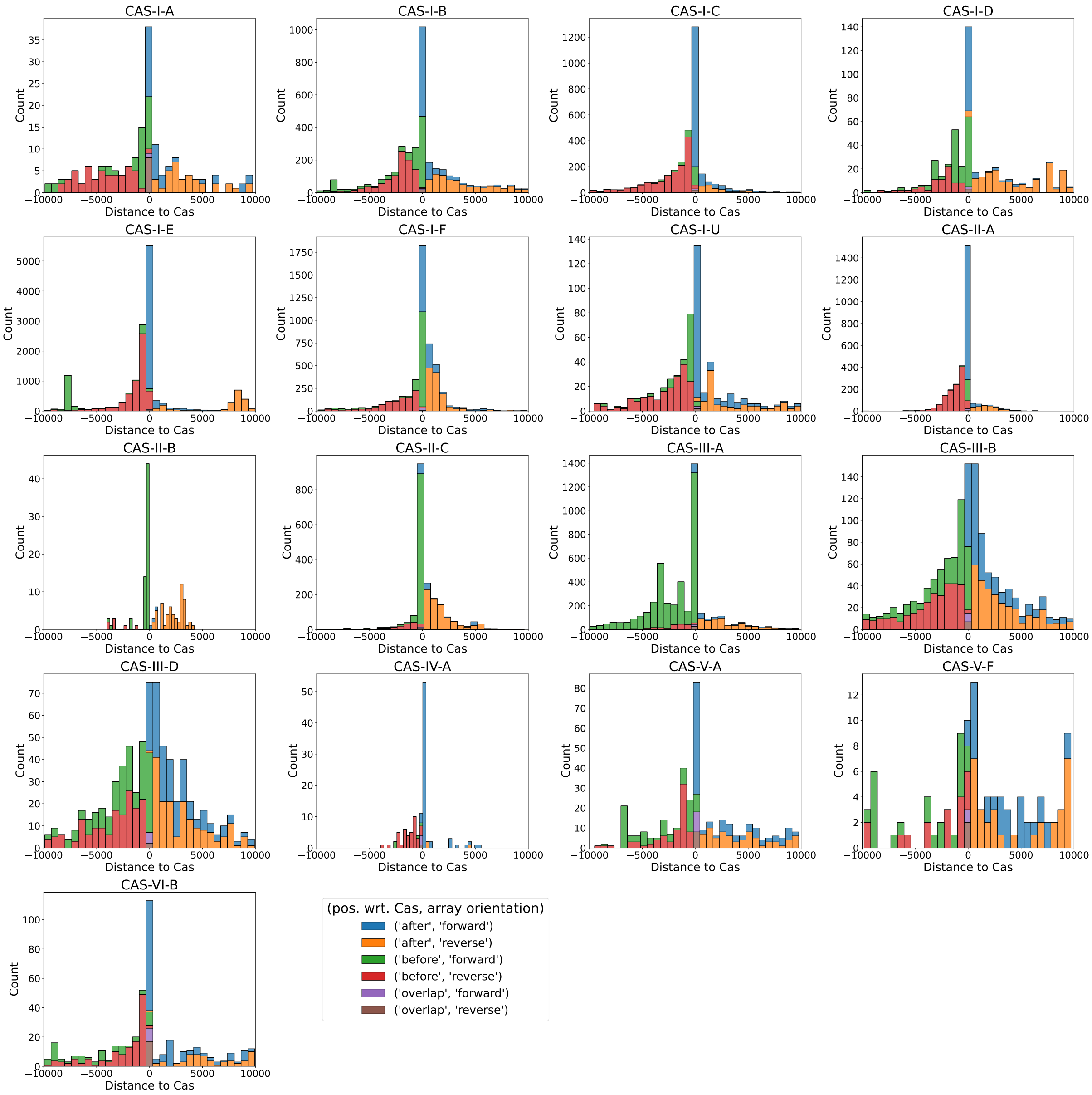


**Supplementary Fig. S8. Directional distances of CRISPR arrays to their respective Cas locus.** We show directional distances from the CRISPR arrays with maximum distance of 10000 bp to their respective Cas locus for all types with sufficient data. The colors show where in the genome the arrays where found and the array orientation. Green and red are 'before' the respective Cas locus (left of (and including) the 0 bin). In blue and orange are arrays 'after' the respective Cas locus (right of (and including) the 0 bin). CRISPR arrays found within the Cas locus are colored in purple and brown and naturally can only be found in the 0 bin. Green, blue and purple mark forward oriented arrays and red, orange and purple mark reverse oriented arrays. Overall, all types favor closeness to the Cas locus but we identify 5 different distribution patterns described in the main manuscript.

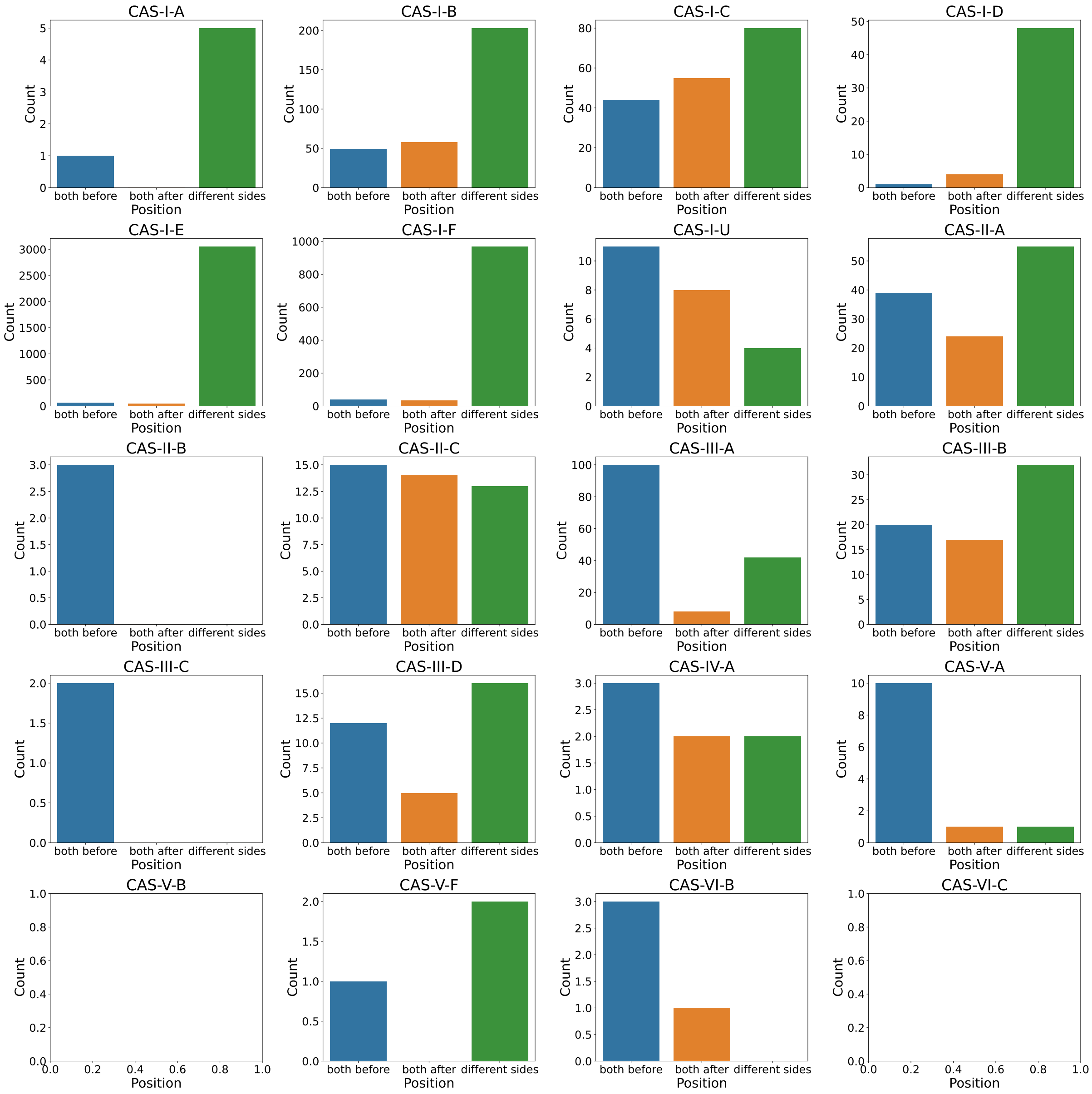


**Supplementary Fig. S9. Positioning of arrays for genomes with exactly two close (< 10000 bp distance) arrays belonging to the same Cas locus.** Type I-D, I-E and I-F greatly favor placing the arrays divided on both sides of the array. Other types are much more diverse in the positioning of the arrays. Note the different y-axis scales for each plot. There is not sufficient data for all types.

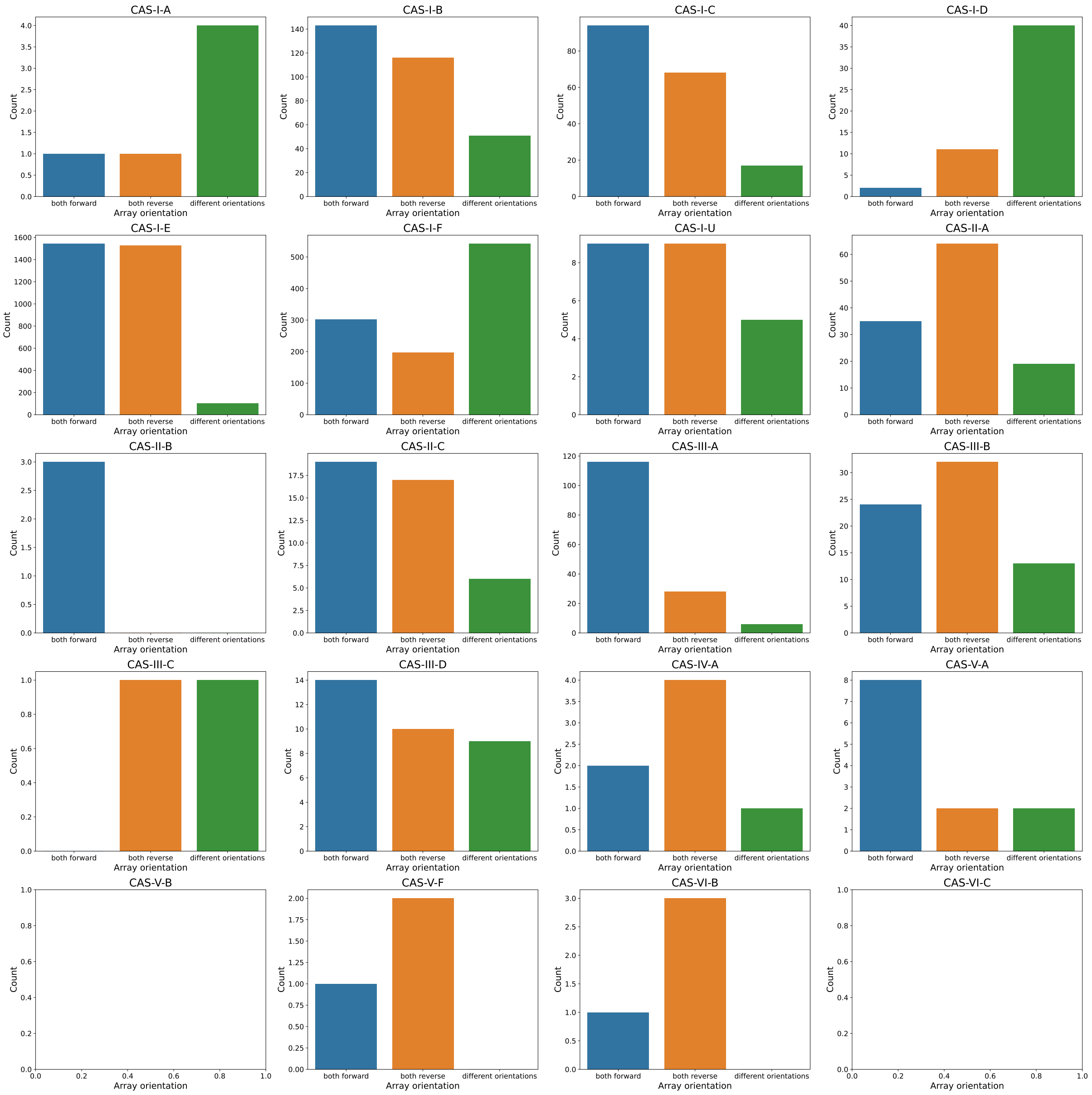


**Supplementary Fig. S10. Orientation of arrays for genomes with exactly two close (< 10000 bp distance) arrays belonging to the same Cas locus.** Note the great differences of distributions of the array orientation between the types I-D, I-E and I-F which often have arrays placed on different sides of the Cas locus, see Figure S9.

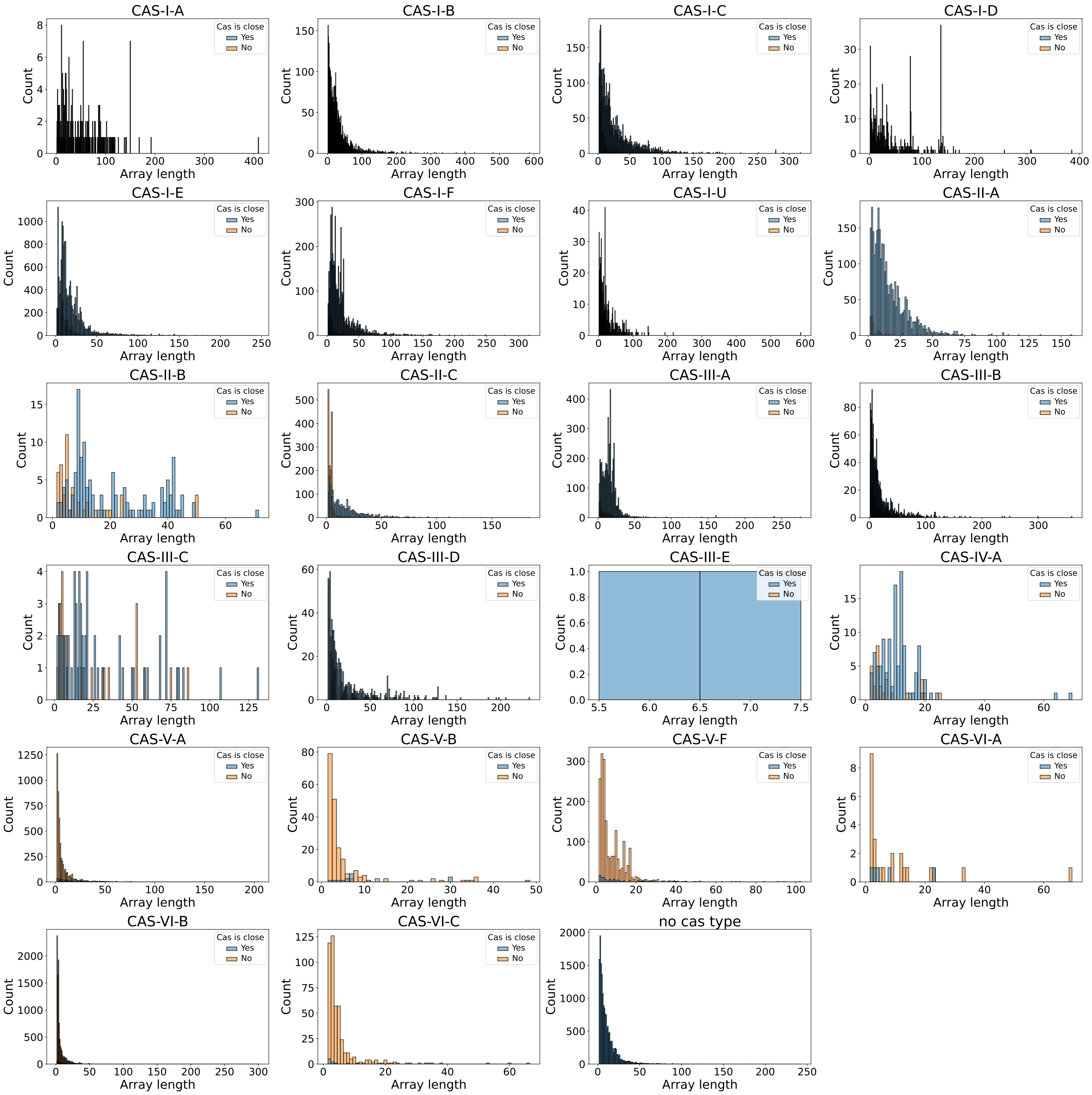


**Supplementary Fig. S11. Spacer array length distributions across Cas types.** We show the distribution of the number of spacers across Cas types with information about the closeness of their Cas locus. The distributions of 'Yes'/'No' for 'Cas is close' are layered over each other. In general, arrays are shorter, if they are further removed from their Cas and there are large differences between types.

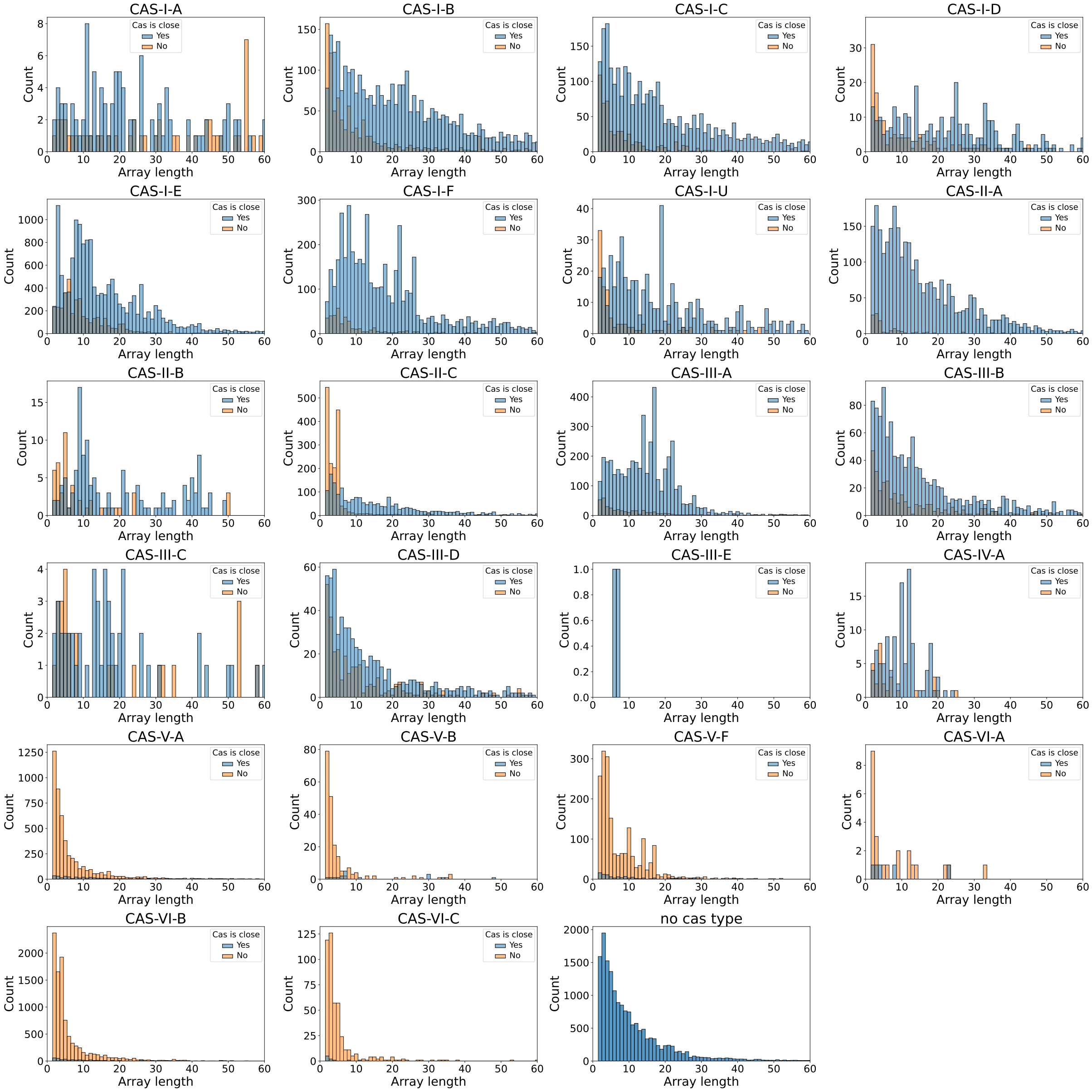


**Supplementary Fig. S12. Spacer array length distributions across Cas types with limited x-axis to 60.** Zoomed in version of the array length distribution plot above.

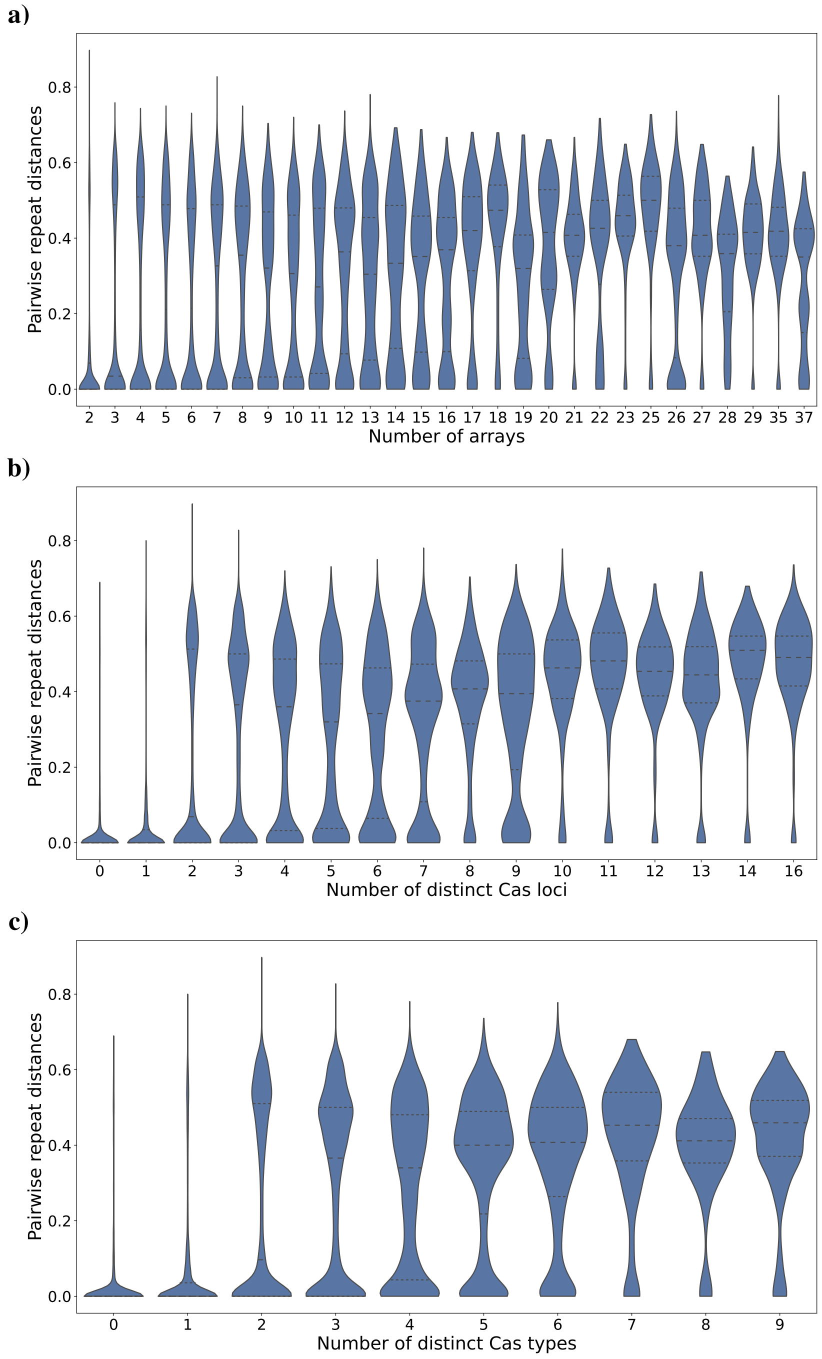


**Supplementary Fig. S13. Distribution of pairwise repeat distances.** We show the distribution of pairwise (normalized) repeat distances with respect to the number of arrays (**a)**), the number of distinct Cas loci within a genome (**b)**) and the number of distinct Cas types within a genome (**c)**). Violins are normalized for each column. It clearly shows that for more arrays or types the distribution becomes bi-or multimodal with peaks around 0, i.e. closely related repeats of interacting arrays (likely of the same Cas type), and higher values (0.4 - 0.5) showing the distances between arrays of (likely) different Cas types. Naturally since we compute pairwise distances the peak in 0 tends to reduce again and genomes carry decent variety of types and repeats leading to multimodal distributions.

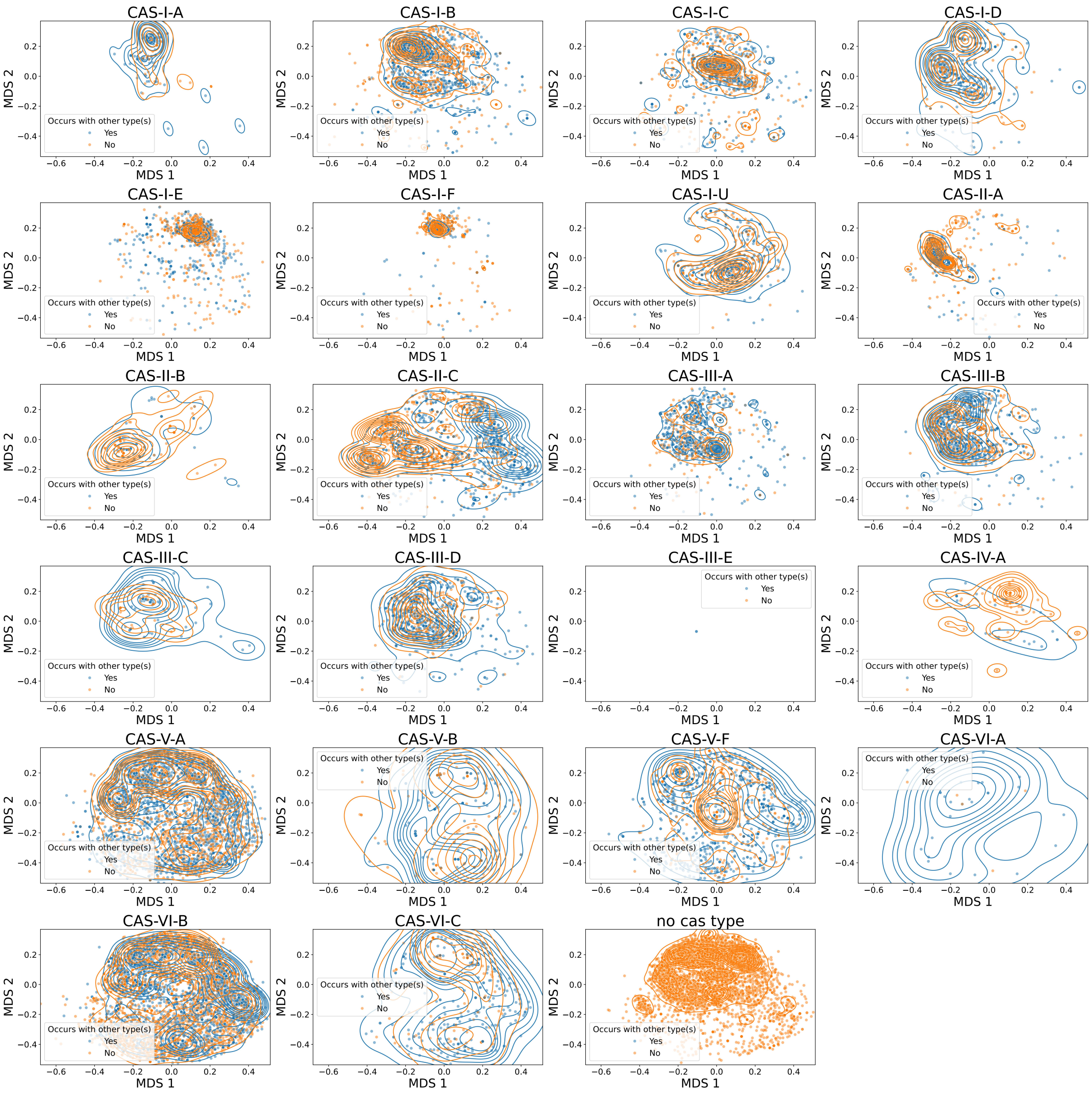


**Supplementary Fig. S14. Repeat MDS into 2-d for all found Cas types (colored by: 'occurs with other type(s)').** Complementary figure to Fig. 4 in the main manuscript showing the repeat MDS for all types in the dataset with respective contours of the kernel density estimates.

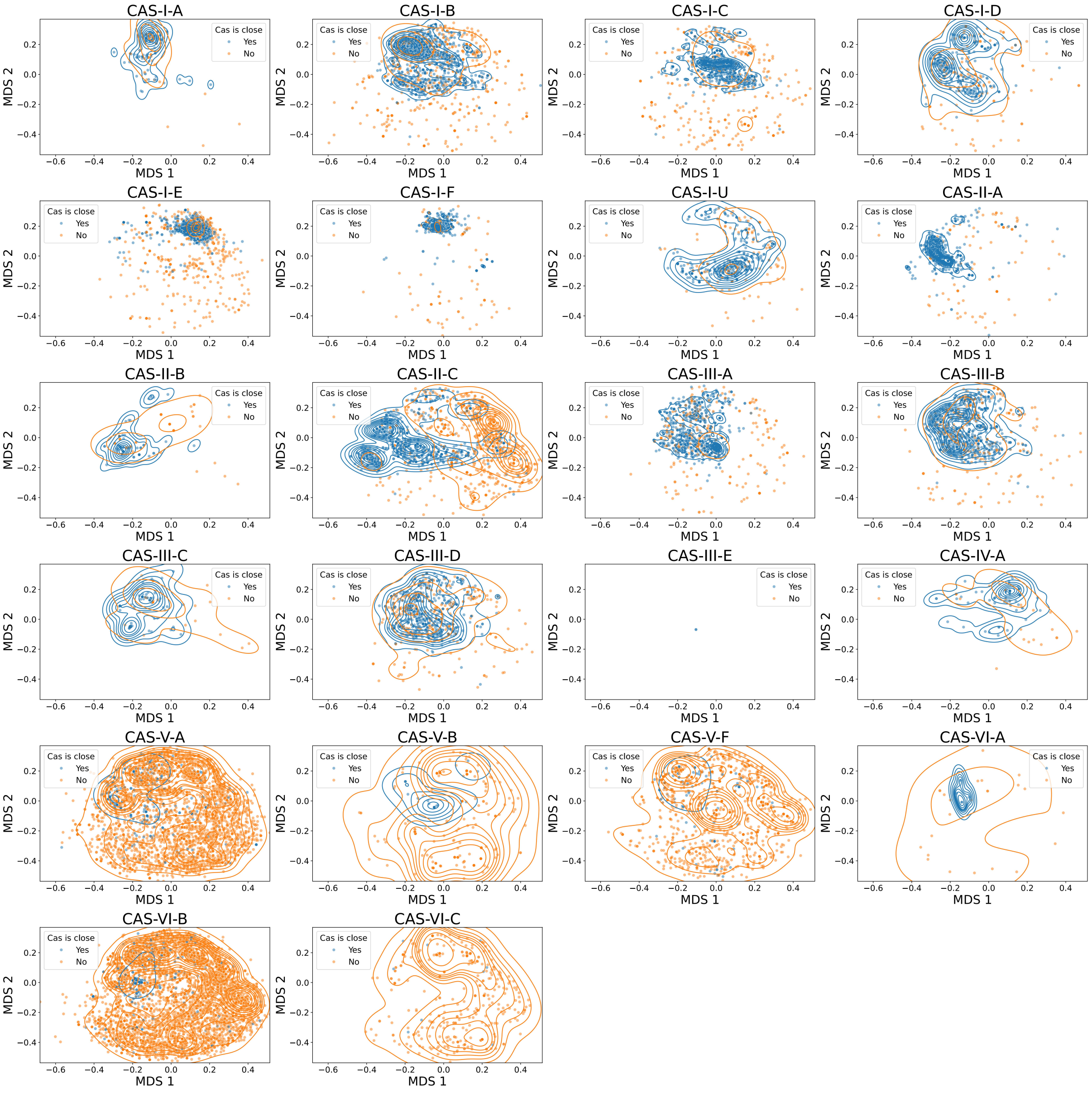


**Supplementary Fig. S15. Repeat MDS into 2-d for all found Cas types (colored by 'Cas is close').** Complementary figure to Fig. 4 in the main manuscript showing the repeat MDS for all types and colored by closeness to the respective Cas locus (close if distance < 10000 bp) with respective contours of the kernel density estimates.
